# The dark transcriptome mouse organogenesis cell atlas

**DOI:** 10.64898/2026.10.06.756869

**Authors:** Miao Liu, Dingyao Zhang, Keyi Li, Yuval Kluger, Jun Lu, Mark B. Gerstein, Haikuo Li, Rong Fan

## Abstract

Single-cell atlases have transformed our understanding of mammalian tissue biology, yet the diverse noncoding “dark transcriptome” remains largely uncharted. Here, we present the Dark Transcriptome Mouse Organogenesis Cell Atlas (DT-MOCA), comprising 276,455 cells across embryonic days 11 to 18 and jointly capturing protein-coding transcripts and five major classes of noncoding RNAs through single-nucleus total RNA sequencing, establishing the first comprehensive catalog of noncoding RNAs across major cell types and developmental lineages. We found that long noncoding RNA and microRNA profiles distinguished major cell types and developmental lineages with resolution comparable to messenger RNA, whereas transfer RNA, small nuclear RNA, and small nucleolar RNA programs were less cell type-discriminative and more broadly shared. We further resolve noncoding RNA expression patterns as lineage-specific, temporally modulated, or coordinately regulated by cell identity and developmental stage. We observed that temporally regulated long noncoding RNAs were evolutionarily younger than those associated with cell identity. We further generate a catalog of the change of noncoding RNAs cross all 11 major developmental trajectories. It revealed lineage-specific coordination between transfer RNA supply and codon demand during musculoskeletal organogenesis as well as dynamic microRNA and small nucleolar RNA programs during the central nervous system development. We observed microRNAs converge on regulators of proliferation and gliogenesis and shared induction of Snord116 defining a developmental window relevant to Prader–Willi syndrome. Together, this atlas provides a comprehensive catalog of the developmental dark transcriptome and reveals how noncoding RNA programs are organized across cell identity, developmental time, and lineage progression.

## Main Text

Protein-coding RNAs account for only ∼2% of all RNA transcripts in mammalian cells, whereas the remaining ∼98% are noncoding RNAs (ncRNAs) that play critical roles across a wide spectrum of cellular functions. These noncoding transcripts comprise a functionally diverse group that includes structural, housekeeping, and regulatory RNAs. Housekeeping ncRNAs, such as transfer RNAs (tRNAs) that decode messenger RNA (mRNA) codons during protein synthesis, small nuclear RNAs (snRNAs) that constitute the catalytic core of the spliceosome, and small nucleolar RNAs (snoRNAs) involved in ribosome biogenesis, are fundamental to core central dogma processes. Meanwhile, regulatory ncRNAs, including microRNAs (miRNAs) that mediate post-transcriptional gene silencing and long noncoding RNAs (lncRNAs) that act via mechanisms ranging from chromatin remodeling to transcriptional scaffolding, orchestrate complex regulatory networks (*1-3*). Collectively, ncRNAs govern essential biological processes spanning cell fate determination, development, and tissue homeostasis (*1-3*). However, conventional single-cell RNA sequencing (scRNA-seq) methods predominantly capture polyadenylated mRNA transcripts, leaving non-polyadenylated ncRNAs largely undetected (*4*). Consequently, single-cell studies have largely been limited to reading mRNA expression, missing the noncoding transcripts that represent the core of RNA-level regulation governing gene activity and cellular function.

To bridge this gap, emerging technologies are beginning to uncover this functionally critical 98% of the "dark transcriptome" in single cells. Recent advances in polyadenylation-based and random primer-based scRNA-seq have begun unlocking the broad spectrum of noncoding transcripts (*5-7*). We recently developed in situ polyadenylation for clinical archival formalin-fixed paraffin-embedded (FFPE) tissues, demonstrating superior performance in spatially co-profiling diverse coding and noncoding species with high coverage (*8*). Together, these emerging technologies began to pave the way for comprehensive profiling of the dark transcriptome, ultimately illuminating the complete RNA regulatory architecture driving tissue development and function.

Mammalian embryogenesis is an exquisitely choreographed process of rapid differentiation, morphogenesis, and spatial organization (*9*). Landmark efforts like the Mouse Organogenesis Cell Atlas (MOCA) transformed our understanding of early development by mapping single-cell lineage trajectories across various stages of early development (*10-12*) and organogenesis (*13-16*). Yet, these foundational atlases remained fundamentally incomplete: because conventional single-cell technologies capture almost exclusively protein-coding mRNAs, the vast noncoding RNA (ncRNA) programs that actively govern gene expression dynamics during development have remained a vast, unexplored terra incognita.

To illuminate this hidden dimension of embryonic developmental biology, we scaled and adapted in situ polyadenylation to create high-throughput single-nucleus total RNA sequencing, leading to the generation of a Dark Transcriptome Mouse Organogenesis Cell Atlas (DT-MOCA). Profiling 276,455 nuclei across embryonic days E11.5–E18.5, DT-MOCA provides a unified, multi-dimensional molecular map simultaneously capturing messenger RNAs alongside all major noncoding RNA classes. This resource reveals previously uncharted, cell-type-specific and temporally dynamic noncoding RNA programs associated with lineage identity and developmental progression, significantly expanding our understanding of the molecular organization of mammalian organogenesis.

### Single-cell total RNA atlas of mouse organogenesis

To construct a large-scale unbiased total RNA transcriptome atlas at single-cell resolution across a number of animals or tissues, we adapted Patho-DBiT (*8*) to develop snTotal-FFPE, a massively parallel split-pool-based single nucleus total RNA sequencing method, to profile both protein-coding and noncoding transcripts in whole mouse embryo FFPE samples. First, we treated FFPE tissue sections with mineral oil for mild deparaffinization and with Tris/2-aminophenylboronic acid hydrochloride for decrosslinking, which demonstrated enhanced recovery of nucleic acids (*17*). Second, after tissue permeabilization, we performed *in situ* polyadenylation by incubating the tissue with poly(A) polymerase, which added artificial poly(A) tails to the 3’ end of RNA transcripts or fragments. Following oligo(dT)-primed reverse transcription, complementary DNA (cDNA) strands of various RNA biotypes were synthesized. Third, we performed three rounds of split-pool barcoding to ligate single-cell barcodes to cDNAs in individual nuclei, followed by library preparation and next-generation sequencing. We applied snTotal-FFPE to mouse embryos collected at 24-h intervals spanning mouse organogenesis from embryonic day 11 (E11) to E18 and constructed a DT-MOCA. This atlas uncovers the total RNA landscape of mouse embryogenesis, highlighting cell-type-specific noncoding RNA dynamics and their roles in lineage development (**Fig. 1A, tables S1-S2**). We profiled a total of 276,455 cells and recovered an average of 1,572 unique molecular identifiers (UMIs) and 700 genes per cell. Compared with recently published mouse embryo atlases (*13, 16*) of protein-coding genes, our study systematically mapped ncRNA expression, capturing 10,752 lncRNAs and 4,192 small ncRNAs across major classes, including miRNAs, snRNAs, snoRNAs and tRNAs. (**Fig. 1B**). As a validation, we aligned reads mapped to the ncRNA genomic loci and verified that most valid reads have non-genomic artificial poly(A) sequences added right next to the 3’ end of mature ncRNA sequences (**Fig. S1A**). Read coverage profiles showed precise coverage of the mature 3p arms of skeletal muscle-enriched miRNAs *Mir1a-1* and *Mir133a-2* and hematopoietic miRNA *Mir142* (**Fig. 1C, Fig. S1A**). Read mappings of the representative tRNA (*tRNA-Asp-GTC*), snRNA (*Rnu1a1*) and snoRNA (*Snord116*) also demonstrated detections of diverse noncoding transcripts (**Fig. S1B**). These analyses benchmarked robust detection of various ncRNAs, including mature miRNAs, tRNAs, snRNAs and snoRNAs, during mouse embryo development.

**Fig. 1.**
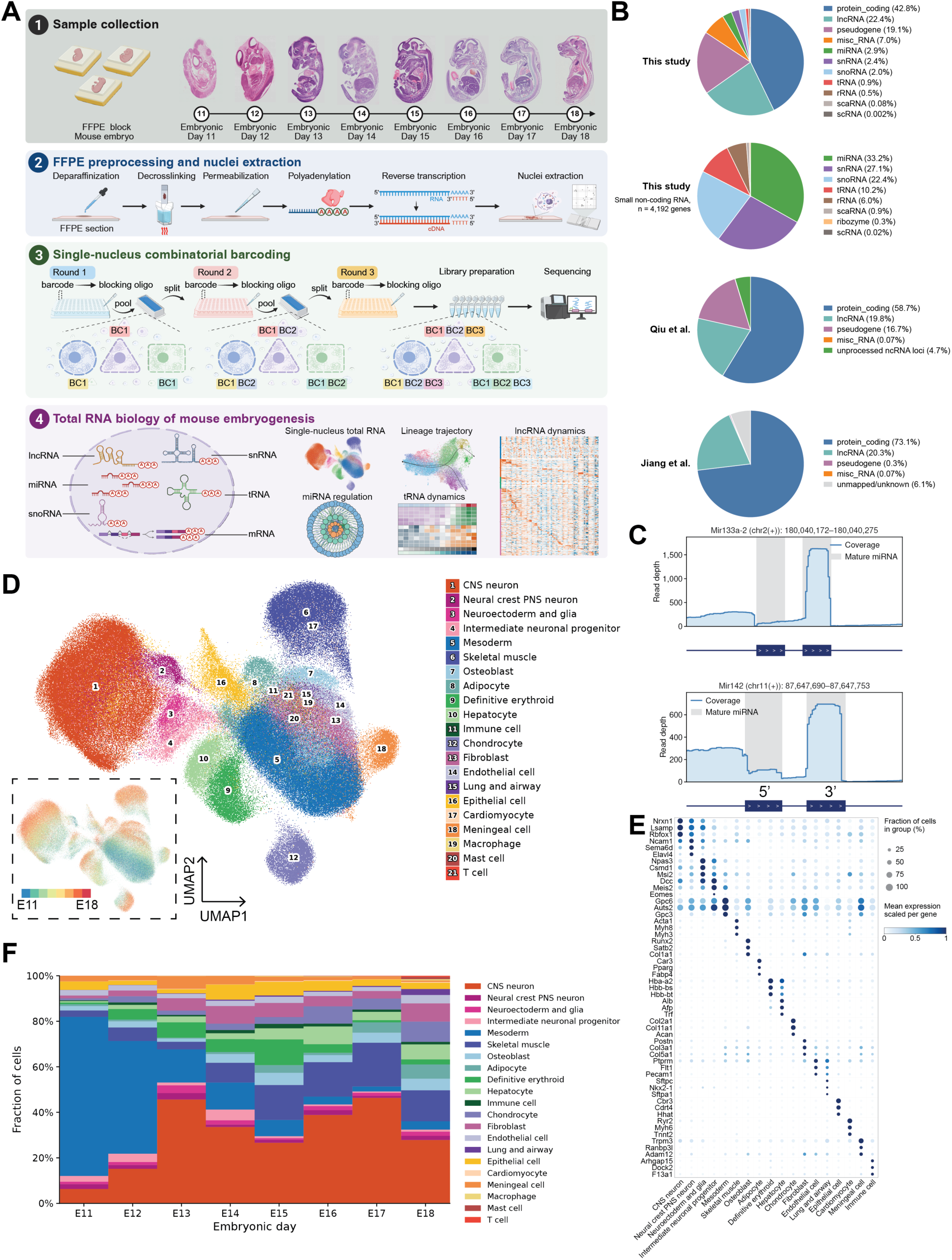
A single-cell temporal atlas of the coding and noncoding transcriptome of mouse organogenesis. **(A)** Schematic illustration of the single-nucleus total RNA sequencing method in formalin-fixed paraffin-embedded (FFPE) mouse embryos spanning embryonic day 11 to day 18. Mouse embryo samples were preprocessed through deparaffinization, decrosslinking, permeabilization, polyadenylation, reverse transcription and nuclei extraction. Single-cell barcodes were added to individual nuclei by three rounds of split-pool combinatorial barcoding. Finally, this atlas revealed total RNA biology of mouse organogenesis, uncovering single-nucleus total RNA expression, lineage trajectory, lncRNA dynamics, miRNA regulation and tRNA dynamics. (**B**) Fraction of gene counts of all RNA biotypes in this study, of small non-coding RNA biotypes in this study, of all RNA biotypes in *Qiu et al.* (*13*), and of all RNA biotypes in *Jiang et al.* (*16*). (**C**) Read coverage for *Mir133a-2* and *Mir142,* with mature miRNA arms shown in gray regions. (**D**) Uniform manifold approximation and projection (UMAP) of single-nucleus total RNA expression profiles, with nuclei colored by cell type. The dashed square showed the same UMAP with nuclei colored by embryonic day. (**E**) Dot plot showing expression of representative protein-coding marker genes that were differentially expressed in each cell type. Dot size indicated the fraction of cells within each cell type with detectable expression. Dot color indicated mean expression across all cells within the cell type, scaled from 0 to 1 for each gene across cell types. Macrophages, mast cells and T cells were grouped within the broad immune cell category. (**F**) Fraction of cell types at each embryonic day. The schematic in (**A**) was created with BioRender.

Unsupervised clustering of single-nucleus total RNA expression identified 21 major cell clusters, annotated based on marker gene expression (**Fig. 1D**, **Fig. 1E, table S3**). We independently validated these annotations by mapping our data to a recent mRNA MOCA of mouse prenatal development spanning late gastrulation to birth (*13*) (**Fig. S1C**). Cell type assignments based on marker gene expression and reference data integration were highly concordant (**Fig. S1D**). Consistent with developmental progression from E11 to E18 (*13*), the proportion of central nervous system (CNS) neurons increased, whereas undifferentiated mesoderm populations declined in relative abundance, accompanied by increased proportions of lineage-restricted derivatives such as skeletal muscle, chondrocyte, and fibroblast cells (**Fig. 1F**). Endoderm-derived hepatocytes also increased in cell type fractions at later stages (**Fig. 1F**). Together, our DT-MOCA atlas captured major embryonic cell types and developmental shifts in cellular composition during mouse organogenesis.

### Cell type-specific ncRNA expression

Genetic and bulk transcriptomic studies have identified regulatory roles for individual ncRNAs in mammalian development, most notably for miRNAs and lncRNAs (*18-20*). However, cell type-specific expression of ncRNAs and their joint regulation across biotypes in individual cells remained unresolved. Mapping ncRNA programs during mouse organogenesis can associate ncRNAs with developmental cell states, nominate candidates for mechanistic investigation, and reveal cell type-restricted ncRNAs with therapeutic potentials. We therefore identified differentially expressed ncRNAs across major cell types within each ncRNA biotype (**table S4**). The lncRNAs detected in DT-MOCA spanned all major genomic classes (**Fig. S2A**) and displayed highly cell type-specific expression patterns (**Fig. 2A**). Cell type-specific lncRNAs included established regulators with versatile functions. Among neuron-enriched lncRNAs, *Paupar* and *Sox2ot* recruited chromatin-modifying complexes to specific locations (*21, 22*). Cardiomyocyte-specific *Mhrt* regulated nucleosome remodeling (*23*), while skeletal muscle-specific *H19* regulated transcription *in cis* at the *Igf2* locus (*24*). Cell type-specific lncRNAs also included miRNA host genes *Mir124-2hg*, *Mir9-3hg*, and *Mir142hg* (**Figs. 2A-2B**). Myogenic *Lincmd1* has been proposed to provide decoy targets for miRNAs (*25*) (**Fig. 2B**). Besides, miRNAs also displayed cell type-specific expression (**Fig. 2C**). Differentially expressed miRNAs included *Mir1a-1* and *Mir133a-2* involved in myoblast differentiation and proliferation (*26*), and *Mir144* required for normal erythroid maturation (*27*) (**Fig. 2D**). Overall, snoRNAs and snRNAs were more broadly expressed across cell types, with snoRNAs showing slightly higher cell type specificity than snRNAs (**Fig. 2E, Fig. S2B**). In particular, *Snord116* family genes were specifically enriched in CNS neurons (**Figs. 2E-2F**). This paternally expressed snoRNA cluster corresponded to a critical noncoding genomic locus implicated in Prader-Willi syndrome (*28*). Prior work documented *Snord116* expression in the embryonic nervous system and neuronal abnormalities following its deletion (*29, 30*). Our atlas extended these findings by resolving cell type specificity of *Snord116* expression at single-cell resolution across E11-E18 organogenesis.

**Fig. 2.**
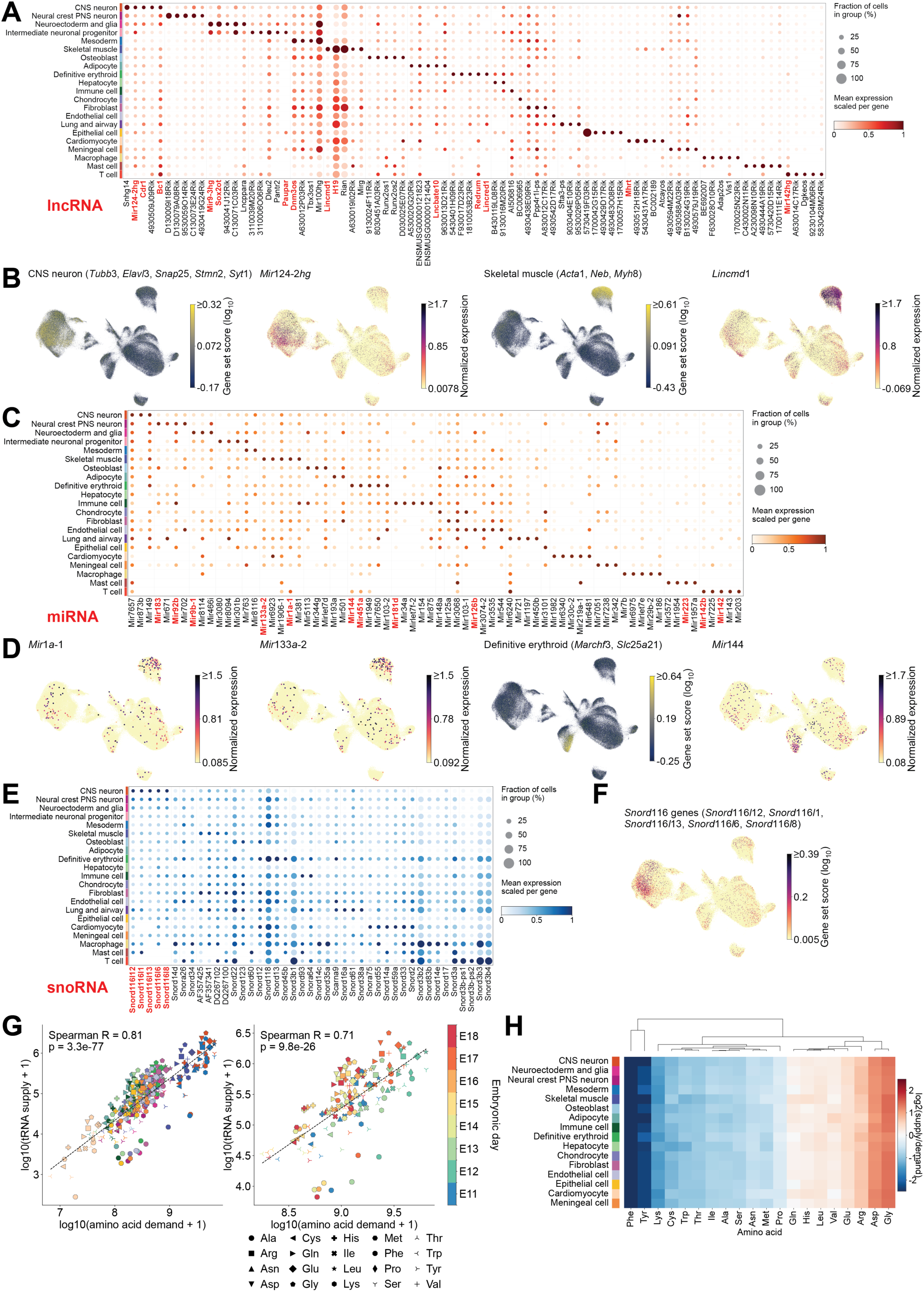
Cell type-specific ncRNA profiles. (**A**) Dot plot showing expression of representative lncRNAs that were differentially expressed in each cell type. The lncRNAs highlighted in red had previously reported functional roles relevant to the corresponding cell types. (**B**) UMAP plots showing CNS neuron (*Tubb3*, *Elavl3*, *Snap25*, *Stmn2*, *Syt1*) and skeletal muscle (*Acta1*, *Neb*, *Myh8*) marker gene scores alongside expression scores for *Mir124-2hg* and *Lincmd1* lncRNAs, highlighting their enrichment in the CNS neuron and skeletal muscle respectively. (**C**) Dot plot showing expression of representative miRNAs that were differentially expressed in each cell type. The miRNAs highlighted in red had previously reported functional roles relevant to the corresponding cell types. (**D**) UMAP plots showing skeletal muscle-enriched *Mir1a-1* and *Mir133a-2* expression, definitive erythroid (*Marchf3*, *Slc25a21*) gene scores, and definitive erythroid-enriched *Mir144* gene expression. Dots with miRNA expression were enlarged in size for visualization. (**E**) Dot plot showing expression of representative snoRNAs that were differentially expressed in each cell type. The snoRNAs highlighted in red had previously reported functional roles relevant to the corresponding cell types. (**F**) UMAP plot showing CNS neuron-enriched *Snord116* gene scores. (**G**) tRNA supply versus amino acid codon demand of each amino acid in each major cell type (left) or embryonic day (right) showed a significantly positive correlation. Spearman correlation coefficients and *P* values from two-sided Spearman rank-correlation tests were shown. Dot color assignment of each cell type (left) is the same as that in panels **A**, **C**, or **E**. (**H**) Heatmap showing the ratio of tRNA supply to amino acid demand across cell types, with amino acids hierarchically clustered using the Euclidean distance. In panels (**A, C, E**), dot size indicated the fraction of cells with detectable expression within each cell type. Dot color indicated mean expression across all cells within the cell type, scaled from 0 to 1 for each gene across cell types.

Furthermore, tRNAs decode mRNA codons and deliver amino acids to the ribosome. The balance between tRNA abundance and the codon composition of mRNAs regulates translational efficiency (*31*). How tRNA abundance or supply varies across cell types during mammalian embryogenesis, and whether it is coordinated with codon demand, has not been resolved at the single-cell resolution. Here, we first identified differentially expressed tRNAs across major cell types, revealing distinct patterns of tRNA isodecoder (defined as tRNAs that carry the same amino acid and share the same anticodon, but differ in sequence elsewhere in the tRNA) enrichment (**Fig. S2C**). For example, CNS neurons were enriched in *tRNA-Arg-TCT-4*, a tRNA known to modulate neuronal translation and synaptic transmission (*32*). Other cell type-associated isodecoders included *tRNA-Ser-AGA-2* in definitive erythroid cells, *tRNA-Tyr-GTA-2* in fibroblasts, and *tRNA-His-GTG-1* in cardiomyocytes (**Fig. S2C**). We further asked whether cell type-specific tRNA programs were adapted to amino acid demands based on codon usage (**Materials and Methods**). We identified a positive correlation between tRNA supply and amino acid demand across cell types (**Fig. 2G**, Spearman R = 0.81, p = 3.3x10^-77^) and across embryonic days (**Fig. 2G**, Spearman R = 0.71, p = 9.8x10^-26^), indicating global alignment between tRNA supply and amino acid demand. However, individual amino acid such as glycine was consistently over-supplied, whereas phenylalanine was consistently under-supplied (**Fig. 2H**). These amino acids also showed consistent supply-demand deviations in human tissues and cell types (*33*), indicating conserved demand mechanisms regulating tRNA supply.

### Temporal dynamics of ncRNA expression

To systematically characterize total RNA expression patterns across developmental stages and cell identities, we modeled the expression of each gene as a function of embryonic day and major cell type using a generalized additive regression model (*34*) (**Fig. 3A**). Within each RNA biotype, we distinguished four gene classes (**Figs. 3A-3B, Figs. S3A-S3B**): genes with shared temporal expression patterns across cell types (Time only), genes differentially expressed across cell types without detectable temporal effect (Cell type only), genes with independent temporal and cell type effects (Time + Cell type, additive), and genes whose temporal trajectories differed by cell types (Time × Cell type, interactive). All four gene classes were resolved in protein-coding genes (**Fig. S3B**), lncRNAs (**Fig. 3B**), miRNAs, tRNAs, snoRNAs and snRNAs. Time-only protein-coding genes that decreased with embryonic days were enriched for ribonucleoprotein complex biogenesis, RNA splicing and tRNA metabolic processes, whereas the three cell type-dependent gene classes were enriched for functions associated with cell identities (**Fig. S3C**). Canonical DNA replication and cell proliferation genes, such as *Mcm5*, *Cul4a*, *Ccnb2*, and *Cenpn*, were also assigned to the Time-only class and declined globally as development proceeded (**Fig. S3D**). These patterns were consistent with bulk transcriptomic studies reporting developmental downregulation of cell cycle, RNA splicing and ncRNA processing genes in mouse embryos (*35*) and declining proliferation genes across developing mouse tissues (*36*). Fractions of the four gene classes varied across ncRNA biotypes (**Fig. 3C**), and directions of developmental changes also differed among them (**Fig. 3D**). The majority of protein-coding genes (67.3%), lncRNAs (54.2%), and snoRNAs (50.5%) decreased during mouse organogenesis, whereas most tRNAs (98.9%) and 48.5% of snRNAs increased (**Fig. 3D**).

**Fig. 3.**
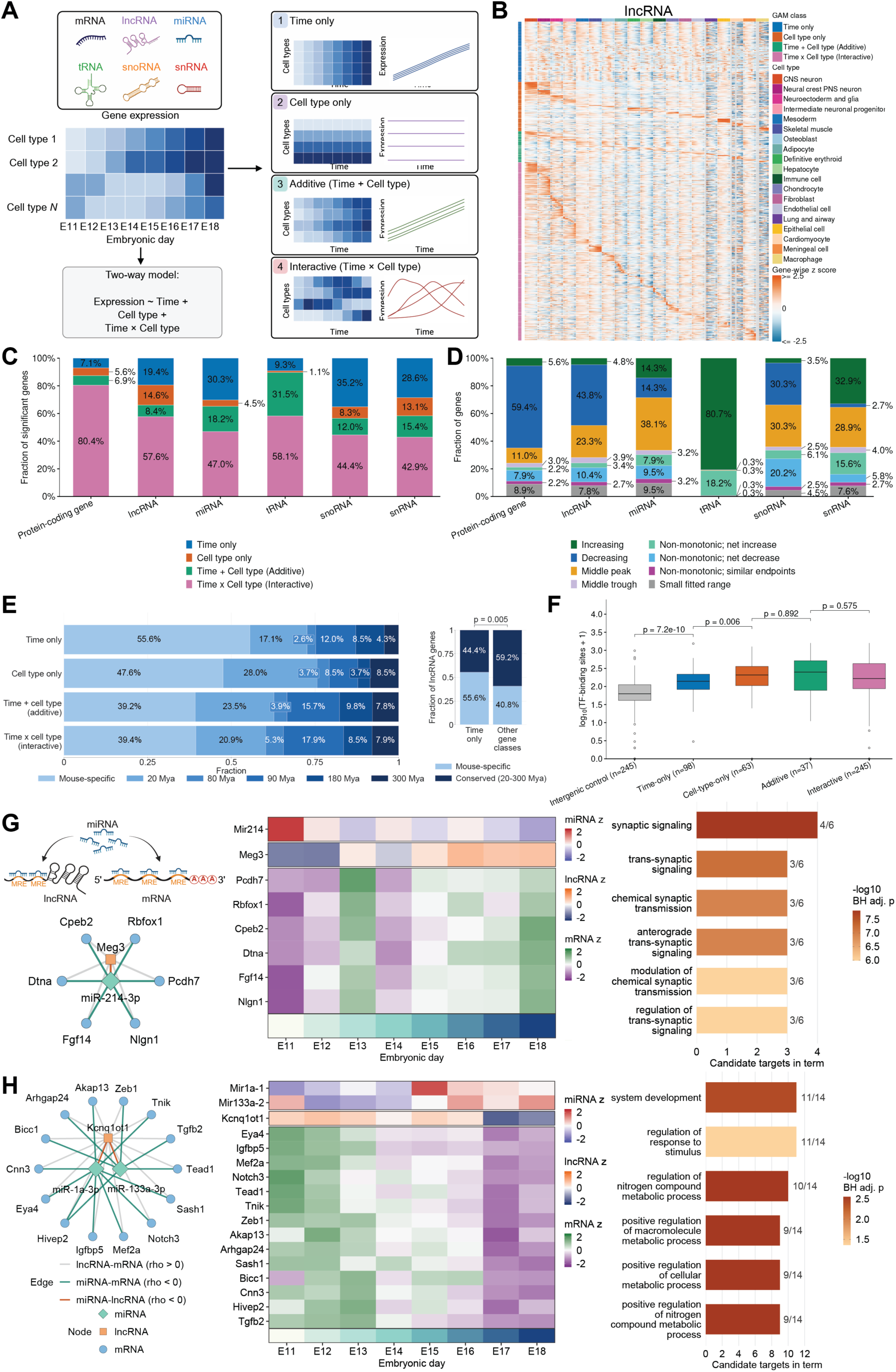
Temporal dynamics of ncRNA expression and regulation. (**A**) Schematic of the generalized additive model framework for modeling gene expression across cell types and embryonic days. Genes within each ncRNA biotype were categorized into four classes based on their modeled expression patterns. (**B**) Heatmap of model-fitted expression of lncRNAs, with rows grouped by the four gene classes and expression pattern modules and columns ordered from E11 to E18 within each major cell type. Colors represented gene-wise z-scores of log2(fitted CPM + 1). CPM: counts per million. Gray indicated cell type–day combinations lacking retained pseudobulk observations. (**C**) Fraction of significant genes assigned to the four gene classes within each RNA biotype of protein-coding gene, lncRNA, miRNA, tRNA, snoRNA and snRNA. (**D**) Fraction of genes with different expression change directions within each RNA biotype of protein-coding gene, lncRNA, miRNA, tRNA, snoRNA and snRNA. (**E**) Fraction of lncRNAs with different evolutionary ages in each lncRNA gene class. The *P* value was calculated from two-sided Fisher’s exact test. (**F**) Distribution of the number of transcription factors predicted to bind to the promoters of lncRNAs of the four gene classes. The *P* values were calculated from two-sided Wilcoxon ranksum tests. (**G**) Left panel: Schematic illustration of the competing endogenous RNA (ceRNA) network. Candidate *Meg3–miR-214-3p* ceRNA network showing positive lncRNA– mRNA Spearman correlations in gray and negative miRNA–lncRNA and miRNA–mRNA correlations in orange and green respectively. Middle panel: Heatmap showing gene-wise z-scores of pseudobulk gene expression for *Mir214*, *Meg3*, and six selected candidate mRNAs across E11– E18. Right panel: Gene ontology bars showing selected biological processes enriched in the corresponding ceRNA module in CNS neurons. (**H**) Left panel: Candidate *Kcnq1ot1–miR-1a-3p/miR-133a-3p* ceRNA network showing positive lncRNA–mRNA Spearman correlations in gray and negative miRNA–lncRNA and miRNA–mRNA correlations in orange and green respectively. Middle panel: Heatmap showing gene-wise z-scores of pseudobulk gene expression for *Mir1a-1*, *Mir133a-2*, *Kcnq1ot1*, and 14 selected candidate mRNAs across E11–E18. Right panel: Gene ontology bars showing selected biological processes enriched in the corresponding ceRNA module in skeletal muscle. The schematic in (**A**) was created with BioRender.

Evolutionary conservation illuminates the functional importance of lncRNAs (*37*). Whether evolutionary age distinguishes temporal from cell type–dependent lncRNA regulation remains unknown. We therefore assigned each lncRNA an evolutionary age using a published dataset (*37*) and compared evolutionary age distributions across the four expression classes. Mouse-specific lncRNAs constituted the largest evolutionary age group in every gene class and accounted for more than half of the Time-only class, whereas conserved lncRNAs were more enriched in the three cell-type-dependent classes (**Fig. 3E, table S5**). To examine transcriptional regulation of the lncRNA gene classes, we compared the number of transcription factors predicted to bind to the lncRNA promoters. Promoters of cell type-dependent lncRNAs were bound by more transcription factors than those of Time-only lncRNAs (**Fig. 3F**), supporting more complex transcriptional regulation. Together, these findings linked lncRNA evolutionary age to distinct modes of developmental regulation. Recently evolved, mouse-specific lncRNAs preferentially participated in temporal programs shared across cell types, whereas evolutionarily older lncRNAs exhibited cell type-dependent expression under more complex transcriptional control.

To associate lncRNAs with biological functions, we constructed gene co-expression networks containing both lncRNAs and mRNAs separately for each cell type, using the hdWGCNA method (*38*). The resulting co-expression modules were enriched for biological processes and signaling pathways relevant to the corresponding cell types (**Fig. S4**). Co-expression with protein-coding programs nominated the lncRNAs as candidates for participation in the corresponding cellular processes, providing a framework for prioritizing functional studies (**Fig. S4**). Some highly abundant lncRNAs can influence post-transcriptional regulation by competing for shared miRNA binding sites (*25*), we thus further constructed candidate lncRNA-miRNA-mRNA competing endogenous RNA (ceRNA) networks (**Materials and Methods**). In CNS neurons, *miR-214-3p* was predicted to target six neuronal differentiation and synaptic genes including *Rbfox1, Cpeb2, Dtna, Fgf14, Nlgn1*, and *Pcdh7*, with the *Meg3* lncRNA predicted to share *miR-214-3p* binding sites, positioning *Meg3* as a candidate regulator of *miR-214-3p* activity (**Fig. 3G**). In skeletal muscle, the myomiRs *miR-1a-3p* and *miR-133a-3p* involved in myoblast proliferation and differentiation were predicted to target muscle-specific transcripts encoding transcription factors *Mef2a*, *Tead1*, *Zeb1* and *Hivep2 (26)* (**Fig. 3H**). The *Kcnq1ot1* lncRNA contained predicted binding sites for both miRNAs, identifying candidate connections between lncRNA expression and miRNA-mediated regulation (**Fig. 3H**). Together, these analyses delineated temporal and cell type-dependent ncRNA programs during mouse organogenesis, resolving lncRNAs underlying evolutionary conservation, transcriptional regulation, and candidate post-transcriptional regulatory networks.

### tRNA dynamics during skeletal lineage specification

We next resolved developmental trajectories that cell types traversed during mouse organogenesis and asked how ncRNAs were dynamically and jointly involved in lineage commitment. We employed Slingshot to infer lineage structures and reconstructed pseudotemporal trajectories of 11 major developmental systems during mouse organogenesis (*39*). Each lineage trajectory was benchmarked by early-stage and late-stage marker gene expression (**Fig. S5, Fig. S6**). Each ncRNA biotype displayed distinct expression dynamics along pseudotime and embryonic days (**Fig. 4, Fig. S7, table S6**). The most complex structures included the musculoskeletal system and neuronal system. The skeletal system included sclerotome, skeletal mesenchymal progenitor, joint interzone, cartilage matrix chondrocyte, prehypertrophic chondrocyte, hypertrophic chondrocyte, osteogenic perichondrial progenitor, preosteoblast, and osteoblast, which recapitulated continuous cascades of chondrocyte and osteoblast maturation (**Fig. 4A**). The muscle trajectory included dermomyotome, myoblast, and myotube (**Fig. 4B**). The neural crest system spanned known developmental trajectories of the peripheral nervous system, from neural crest sensory progenitors and Schwann cell precursors through immature sensory neuroblasts to differentiated large-diameter and small-diameter sensory neurons, as well as sympathetic noradrenergic and enteric neurons (**Fig. 4C, Fig. S7A**). The dorsal hindbrain/spinal neuronal system included dorsal neural progenitor, dorsal somatosensory relay neuron, glycinergic inhibitory spinal interneuron, Group-Z spinal neuron, maturing hindbrain/spinal neuron, and immature hindbrain/spinal neuron (**Fig. 4D, Fig. S7B**). The cortical neuronal trajectory included radial glia, early postmitotic cortical neuron, cortical projection neuron, and Cajal-Retzius cells (**Fig. 4E**). The hematopoietic trajectory included hematopoietic stem and progenitor cell, proliferating erythroblast, terminal erythroblast and erythroid (**Fig. 4F**). We further delineated fibroblast, meningeal, smooth muscle, hepatic and endothelial trajectories spanning progenitor and more differentiated cell states (**Figs. 4G-4K, Fig. S7C**).

**Fig. 4.**
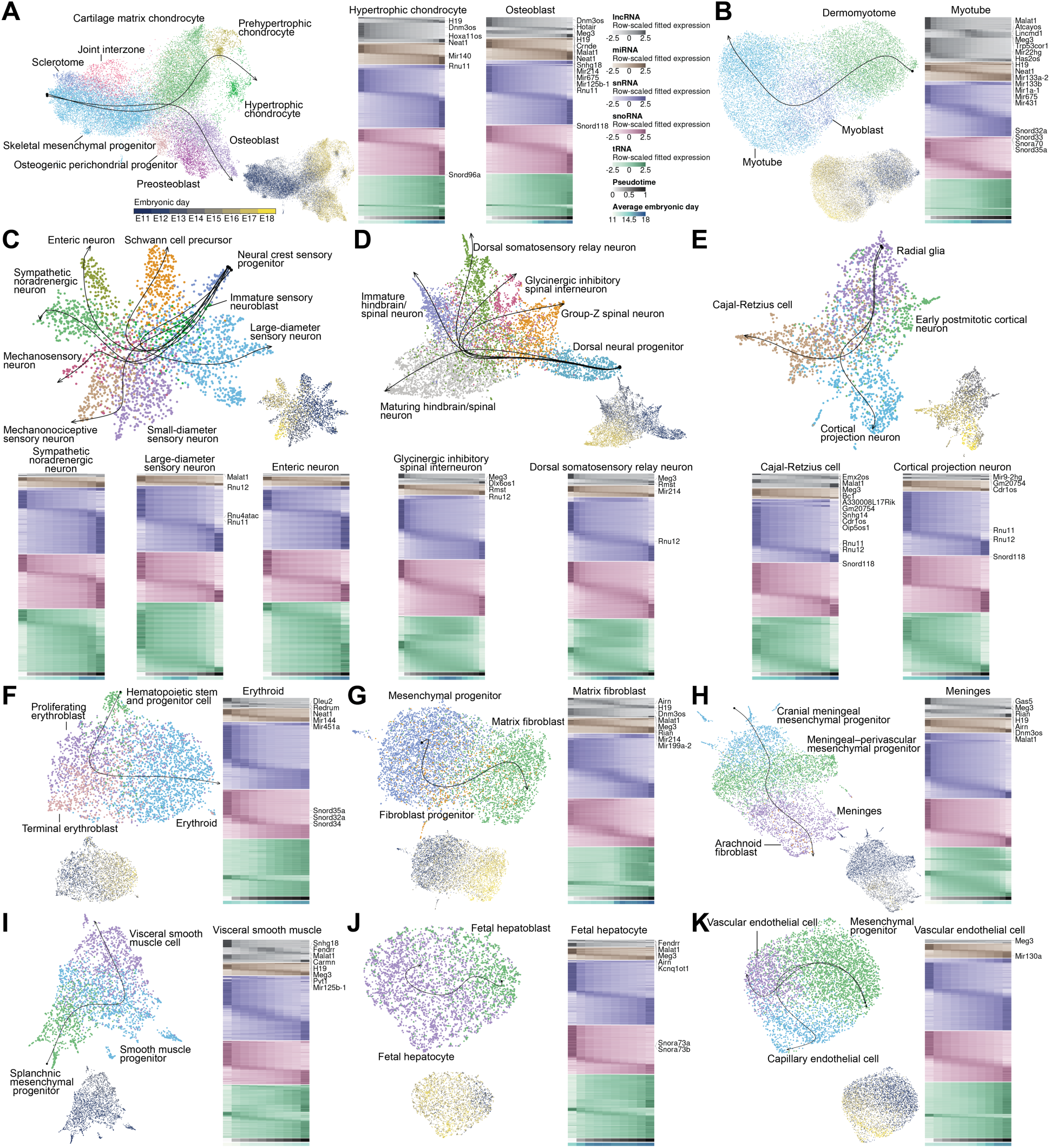
Characterization of ncRNA dynamics along 11 major developmental trajectories. UMAP of single-nucleus total RNA expression, with nuclei colored by cell types during the development of the skeletal system (**A**), skeletal muscle (**B**), neural crest (**C**), dorsal hindbrain/spinal neuronal system (**D**), cortical neuronal system (**E**), hematopoiesis (**F**), fibroblast (**G**), meningeal cell (**H**), smooth muscle (**I**), fetal hepatocyte (**J**), and endothelial cell (**K**). Developmental trajectories fitted from Slingshot were projected onto the UMAP plots. The heatmap showed expression dynamics for lncRNAs, miRNAs, snRNAs, snoRNAs, and tRNAs along each developmental trajectory, with rows grouped by RNA biotype and temporal expression patterns and columns representing pseudotime bins. Colors indicated gene-wise z-scores of gene expression. Bottom bars indicated pseudotime and mean embryonic day.

Much of the mammalian skeleton forms through endochondral ossification, which coordinates chondrocyte maturation with osteoblast-mediated bone formation (*40*). Collagens constitute the principal structural components of the organic matrix in bone and cartilage. The defining feature of collagen is a repeating GlyXaaYaa sequence where every third residue is a glycine (*41*). Chondrocytes and osteoblasts therefore sustain an exceptional glycine demand as they build the collagenous matrix of the embryonic skeleton. However, it remains unknown how glycine tRNAs change during chondrogenic and osteogenic lineage specification. We found that the three glycine tRNA isoacceptors (defined as tRNAs that carry the same amino acid but have different anticodons, *Gly-GCC*, *Gly-TCC*, *Gly-CCC*) were highly expressed in chondrocytes and osteoblasts, whereas their expression levels were lower in skeletal mesenchymal progenitors (**Fig. 5A, Fig. S5A**). *Gly-GCC* transcripts accounted for the largest fraction of the glycine tRNA pool in both lineages (**Fig. 5B**), consistent with the predominance of the *tRNA-Gly-GCC* isoacceptor reported in human HEK293 cells (*42*). To further quantify how glycine tRNAs are coordinated with codon demand changes along chondrogenic and osteogenic differentiation trajectories, we temporally tracked glycine codon demand (expression-weighted codon frequency across protein-coding sequences) and tRNA supply (tRNA abundance normalized to total non-mitochondrial tRNA counts) changes (**Fig. 5C**). Glycine codon demand increased along both chondrogenic and osteogenic trajectories, accompanying collagen gene upregulation (**Fig. 5D**). Notably, all four glycine tRNA isoacceptors showed an overall increase during differentiation (**Fig. 5D**). Pseudotime profiles indicated that the rise in glycine tRNA abundance preceded the increase in glycine codon demand in both lineages (**Fig. 5E**). Mapping these profiles to embryonic days placed the earliest detectable tRNA increase near E12 and demand onset near E15, with both measures reaching their highest observed levels in late trajectory states around E16 (**Fig. 5E**). Developmental upregulation of glycine tRNAs has also been reported during human mesenchymal stem cell chondrogenesis *in vitro* (*43*). To explore transcriptional mechanisms of glycine tRNA upregulation in the chondrogenic lineage, we examined transcripts encoding RNA polymerase III subunits and TFIIIB/TFIIIC initiation factors, together with glycine pre-tRNA junction signals (**Figs. S8B-S8C**). TFIIIB/TFIIIC transcripts were relatively enriched in early development, whereas Pol III subunit expression varied modestly (**Figs. S8B-S8C**). Glycine pre-tRNA junction signals peaked transiently near E15, whereas mature glycine tRNA abundance reached its maximum around E16 and remained elevated in late developmental states (**Figs. S8B-S8C**). Together, these profiles revealed an earlier increase in glycine tRNA abundance than codon demand, accompanied by distinct precursor and mature tRNA dynamics during skeletal differentiation (**Fig. S8D**).

**Fig. 5.**
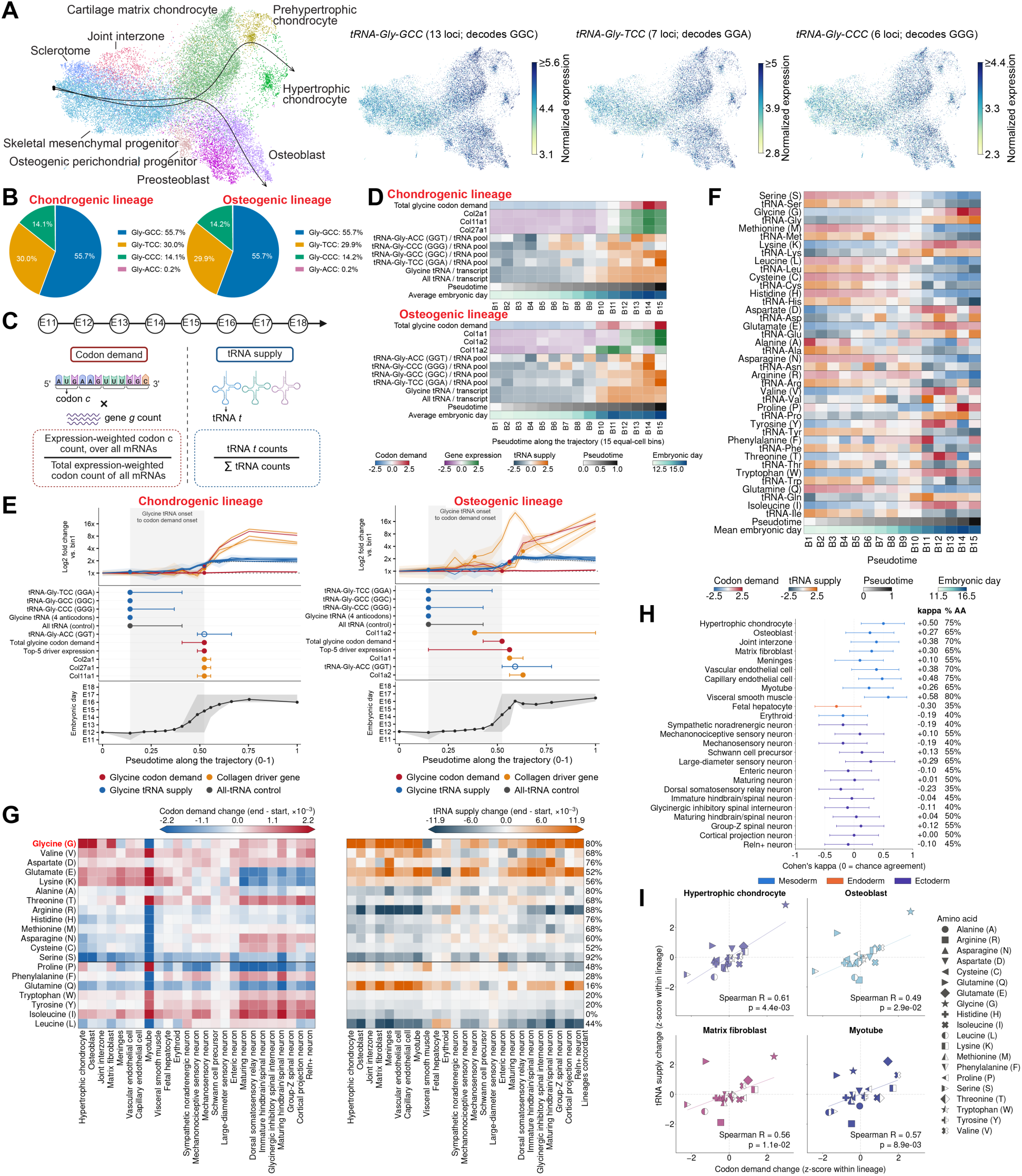
Lineage-specific coordination of tRNA supply and codon demand during mouse musculoskeletal organogenesis. (**A**) UMAP of single-nucleus total RNA expression, with nuclei colored by cell types during the development of the skeletal system. Developmental trajectories fitted from Slingshot were projected onto UMAP. UMAP plots showing *tRNA-Gly-GCC*, *tRNA-Gly-TCC* and *tRNA-Gly-CCC* were shown on the right. (**B**) Proportions of glycine tRNA UMI counts contributed by each anticodon class in the chondrogenic and osteogenic lineage. (**C**) Schematic illustration of the codon demand and tRNA supply calculation. (**D**) Heatmaps showing total glycine codon demand, collagen gene expression, and glycine tRNA abundance across equal-cell pseudotime bins along chondrogenic (upper) and osteogenic (lower) lineages. Anticodon-specific tRNA counts were normalized to total tRNA expression, whereas aggregate glycine tRNA abundance was normalized to total UMI counts. Colors represented row-wise gene expression z-scores, with pseudotime and average embryonic day indicated below. (**E**) Onset ordering of glycine tRNA abundance, glycine codon demand, and collagen expression along chondrogenic and osteogenic developmental trajectories. The top panel showed log2 gene expression fold changes of 15 pseudotime bins relative to the first bin. The middle panel showed onset estimates with 95% bootstrap intervals from 4,000 within-bin cell resamples. Onset was defined as the first bin whose lower 95% confidence limit exceeded the baseline upper limit and remained above it thereafter. The gray band spanned the glycine tRNA-to-demand onset interval. Open symbols indicated low-abundance *tRNA-Gly-ACC* (<1% of glycine tRNA UMIs). The bottom panel showed mean embryonic day per bin, with shading indicating the interquartile range. (**F**) Heatmap showing codon demand and tRNA supply for 20 canonical amino acids across equal-cell pseudotime bins along the chondrogenic lineage. (**G**) Late-minus-early changes in amino acid codon demand and tRNA supply for 20 amino acids across 25 lineages. Percentages indicated the proportion of lineages with concordant directions of change for each amino acid. The divider separated amino acids with ≥50% versus <50% directional agreement. (**H**) Cohen’s kappa quantified the agreement between codon demand and tRNA supply change directions beyond chance. Points and horizontal bars indicated kappa estimates and 95% bootstrap intervals. Percentages reported amino acids with matching change directions. (**I**) Late-minus-early changes in tRNA supply versus codon demand showed positive correlations across 20 amino acids in four selected lineages. Changes were z-scored within each lineage. Spearman correlation coefficients and *P* values from two-sided Spearman rank-correlation tests were shown. The schematic in (**C**) was created with BioRender.

We next asked whether changes in tRNA abundance were coordinated with codon demand across the 20 canonical amino acids. Within the chondrogenic lineage, tRNA supply and amino acid codon demand changed in the same direction for 15 of 20 amino acids (**Fig. 5F**). Across 25 developmental trajectories, 13 amino acids showed concordant changes in at least half of the lineages (**Fig. 5G**). Directional agreement was the strongest in mesoderm lineage trajectories such as chondrogenic, endothelial and smooth muscle trajectories and generally weaker in neuronal trajectories (**Fig. 5H**). Across amino acids, endpoint changes in tRNA supply and amino acid demand were positively correlated in chondrogenic, osteogenic, matrix fibroblast, and myotube trajectories (**Fig. 5I**). Together, these findings revealed that developmental remodeling of tRNA supply aligned with changing codon demand in a lineage- and amino acid-dependent manner, with stronger tRNA supply-codon demand coordination during musculoskeletal organogenesis.

### ncRNA landscape of the developing mouse brain

During mouse CNS development, radial glia generally progressed from neurogenesis towards gliogenesis, with regional differences in timing and astrocyte production extending from prenatal into postnatal stages (*44*). However, it remained unexplored which ncRNA programs distinguished these developmental trajectories. We thus reconstructed developmental trajectories delineating lineage progression from radial glia to maturing postmitotic neurons and to astrocyte precursors (**Fig. 6A**). Cell identities were supported by expression of neural progenitor regulators *Gli3* and *Prdm16* in radial glia, synaptic genes *Syt1* and *Celf4* in maturing postmitotic neurons, and astrocytic transporter genes *Slc6a11* and *Slc4a4* in astrocyte precursors (**Fig. 6B**). Because miRNAs fine-tune the transcriptome by repressing networks of protein-coding genes, we next identified candidate post-transcriptional programs accompanying lineage divergence (**Fig. 6B**). In particular, *Mir770*, *Mir1906-1*, *Mir7657, Mir6516*, *Mir690* and *Mir485* increased specifically in the neurogenic trajectory. We integrated target predictions using multiMiR (*45*), ENCORI (*46*) and DIANA-LncBase v2 (*47*), retaining miRNA-target pairs co-detected in the same cells. Inverse expression relationships linked *Mir770* to the proliferation and progenitor-associated genes *Hmga2*, *Ttyh1*, and *Cdk6*; *Mir1906-1* to *Nfia*, *Zeb1*, and *Ptprz1*; *Mir7657*, *Mir6516*, and *Mir690* to *Nfia* and *Nfib*; and *Mir485* to *Npas3*, *Nfib*, and *Dlc1* (**Fig. 6C**). The convergence of several miRNAs on *Nfia* and *Nfib*, transcription factors implicated in gliogenic competence (*48*), suggested a candidate network that could reinforce neuronal commitment by attenuating progenitor maintenance and alternative glial programs. Predicted interactions also connected *Mir770* with *Mir9-3hg*, *Mir1906-1* and *Mir6516* with *2610307P16Rik*, and *Mir6516* with *2610035D17Rik*, extending these candidate networks to lncRNAs.

**Fig. 6.**
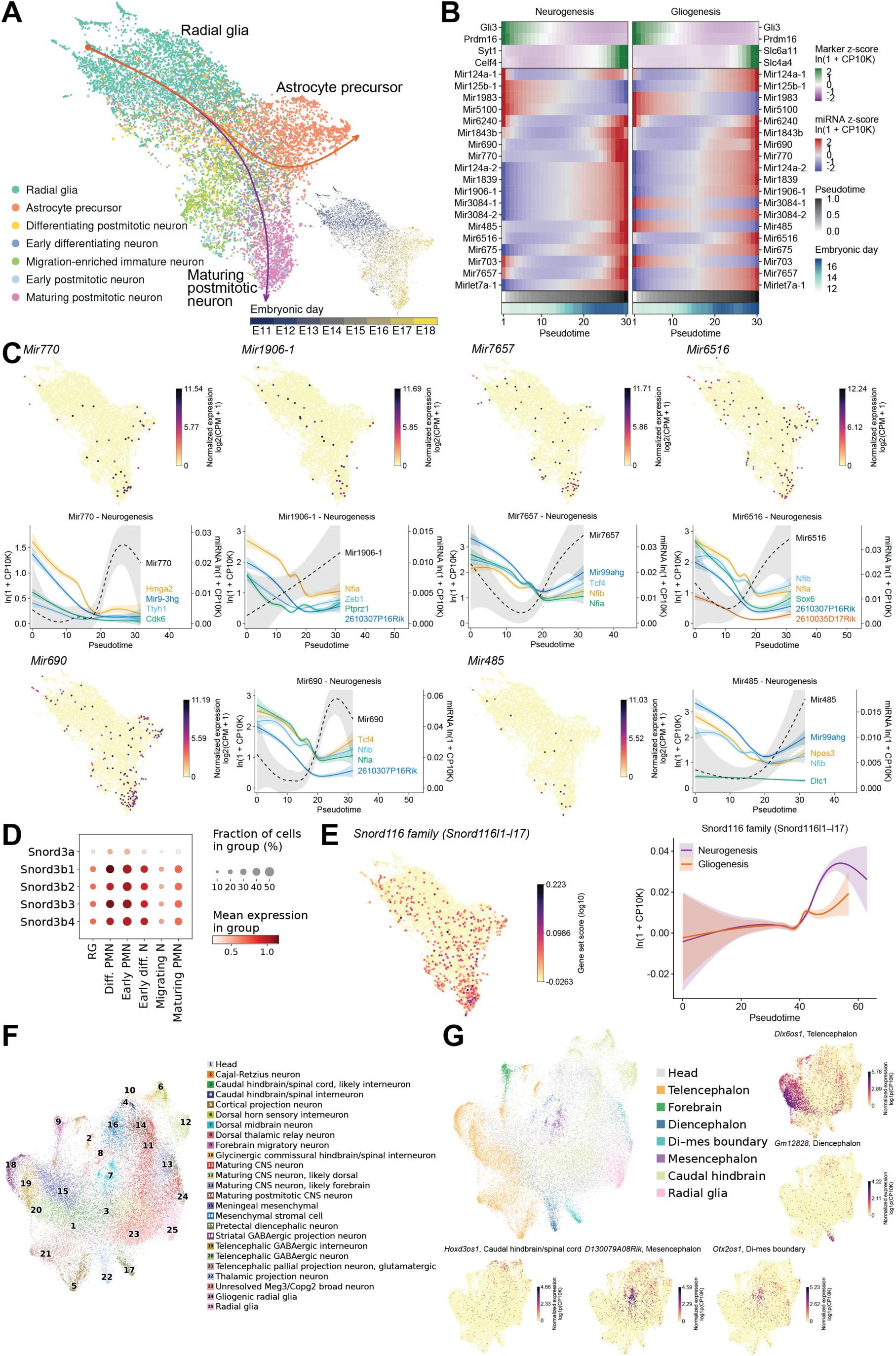
Dynamics of ncRNA profiles and regulation in the developing mouse brain. (**A**) UMAP of single-nucleus total RNA expression from the developing mouse CNS, with nuclei colored by cell types. The same UMAP showing nuclei colored by embryonic days was displayed on the lower right. (**B**) Heatmaps showing early-stage and late-stage marker genes and selected miRNAs along neurogenic and gliogenic trajectories. Colors indicated gene-wise z-scores within each lineage. (**C**) UMAP plots of miRNA expression. Dots with miRNA expression were enlarged in size for visualization. In (**B**) and the expression curves in (**C**), gene expression was calculated as ln(1+10,000×gene UMIs/total cellular UMIs). Expression curves for miRNAs and predicted target genes along the neurogenic trajectory were fitted using the geom_smooth function in ggplot2, with shaded bands indicating 95% confidence intervals. (**D**) Dot plot showing expression of *Snord3a* and *Snord3b1-4* snoRNAs which were differentially expressed in each of the radial glia (RG), differentiating postmitotic neuron (Diff. PMN), early postmitotic neuron (Early PMN), early differentiating neuron (Early diff. N), migration-enriched immature neuron (Migrating N), and maturing postmitotic neuron (Maturing PMN). (**E**) UMAP plot showing *Snord116* family (*Snord116l1*–*Snord116l17*) gene expression. The curve plot showed mean expression of *Snord116* family genes along neurogenic and gliogenic trajectories, fitted using the geom_smooth function with shaded bands indicating 95% confidence intervals. Gene expression was calculated per nucleus by averaging gene-level ln(1+10,000×gene UMIs/total cellular UMIs) values. (**F**) UMAP of single-nucleus total RNA expression in CNS neurons, with nuclei colored by cell types. (**G**) The same UMAP as (**F**) but with nuclei colored by mouse brain regions along the anterior-posterior axis. UMAP plots showing region-enriched lncRNAs were also displayed.

SnoRNA dynamics revealed distinct programs associated with neuronal differentiation. U3 snoRNAs encoded by *Snord3a* and *Snord3b1*-*Snord3b4*, which participated in pre-rRNA processing and 18S rRNA maturation (*49*), were transiently elevated in early differentiating, early postmitotic, and differentiating postmitotic neurons (**Fig. 6D**). This pattern suggested a discrete window of elevated ribosome biogenesis during early neuronal differentiation. Meanwhile, *Snord116* family genes increased specifically along the neurogenic trajectory (**Fig. 6E**). *Snord116* is a paternally expressed, imprinted snoRNA cluster whose deletions encompassing the human *SNORD116* region caused Prader–Willi syndrome (*50*). Consistent with reported *SNORD116* accumulation during neuronal differentiation in human cell culture (*51*), our analysis resolved its preferential increase along embryonic neuronal rather than astrocyte lineage differentiation, providing a developmental context for investigating the consequences of its loss.

To determine how these lineage-associated ncRNA programs relate to diverse neuronal cell types and brain regions, we performed unsupervised clustering of single-cell total RNA expression in CNS neurons and annotated clusters by cell identities and anterior-posterior positions (**Figs. 6F-6G**). Regionally enriched lncRNAs delineated spatially restricted gene expression programs (**Fig. 6G**), including *Dlx6os1* in telencephalon, *Gm12828* in diencephalon, *Otx2os1* in diencephalon-mesencephalon boundary, *D130079A08Rik* in mesencephalon, *Hoxd3os1* in caudal hindbrain/spinal cord. Next, to distinguish ncRNA programs associated with neuronal identities from those with developmental timing, we used generalized additive regression models across cell type, anterior–posterior, dorsal–ventral, and excitatory–inhibitory annotations (**Fig. S9A**). Most *Snord116* members belonged to the Time-only class, with significant enrichment in the cell type and anatomical analyses but no enrichment in whole embryo or neural crest/PNS analyses (**Fig. S9B**). Both *Snord116* and the neighboring *Snord115* gene cluster showed their highest expression in CNS neurons (**Figs. S9C-S9D**). Across CNS groups, *Snord116* exhibited a common rise around E15 across anterior–posterior, dorsal–ventral, and excitatory–inhibitory identities (**Fig. S9E**), indicating its shared temporal program across CNS neuronal identities. Temporal profiles placed the half-maximal rise of *Rbfox3* near E12 and its peak at E13, preceding *Snord116* and *Snord115* half-maximal rises around E13.5 and maxima at E15–E16 (**Fig. S9F-S9G**). *Snrpn* reached its highest sampled expression at E18 (**Fig. S9F-S9G**). The earlier *Rbfox3* increase was consistent with its reported role in promoting neuronal splicing of the *Snord116* host transcripts (*52*). Given the association of paternal human *SNORD116* deletions with the Prader-Willi syndrome (*50*), this shared embryonic expression program provided a temporal framework for investigating the disease mechanisms and nominated E15–E16 as a window for testing its contributions to neuronal maturation.

## Discussion

In this work, we constructed DT-MOCA, a single-cell temporal atlas of the coding and noncoding transcriptome during mouse organogenesis. We profiled 276,455 cells from mouse embryos spanning E11 to E18, resolving the temporal dynamics of lncRNA, miRNA, tRNA, snRNA, and snoRNA expression at single-cell resolution. The atlas was constructed by a snTotal-FFPE method that can map total RNA expression in single nuclei in FFPE tissues. Although mature RNAs are predominantly cytoplasmic, it has been known that nuclear RNA profiles broadly reflect their relative abundance in whole cells. Residual cytoplasmic material may also contribute to detection, although the RNA regulation inferred from nuclear data should be interpreted cautiously. snTotal-FFPE supports multiplexing multiple embryos per experiment, costs $0.01 per cell which is comparable to standard scRNA-seq (*53*), and uses standard laboratory equipment, facilitating adoption by individual laboratories. Beyond developmental studies, snTotal-FFPE can construct total RNA atlases from archived clinical specimens and further enable single-cell profiling of multiple modalities in complex tissues.

DT-MOCA has revealed ncRNA dynamics across embryonic days, cell types and developmental lineages. Globally, lncRNAs with shared temporal expression patterns across cell types were more mouse-specific and newly evolved, whereas lncRNAs associated with cell type identities were more evolutionarily conserved. This functional distinction was corroborated by more predicted transcription factor binding sites on promoters of lncRNAs associated with cell types, which implicated more complex transcriptional regulation. Specifically, we identified cell type-specific lncRNAs and lncRNA-mRNA co-expression modules and nominated lncRNA-miRNA-mRNA competing endogenous RNA networks that can regulate gene activities post-transcriptionally. We further constructed lineage developmental trajectories and identified dynamic ncRNA expression programs along developmental trajectories.

In particular, how differentiating cells coordinate their tRNA abundance with changing amino acid demand remains incompletely understood. Proliferation-specific and differentiation-specific tRNAs typically carried anticodons that matched the codon usage of cell-autonomous and multicellularity genes respectively (*54*). A recent work further showed that tRNA anticodon pools were more stable among cell types despite extensive remodeling of individual tRNA transcripts during *in vitro* differentiation of human induced pluripotent stem cells into neuronal and cardiac cells (*55*). These results suggested that cell differentiation can be supported by both selective tRNA transcript remodeling and maintenance of an existing tRNA anticodon decoding repertoire. In our study, skeletal differentiation was characterized by collagen induction and increased glycine demand, reflecting the requirement for glycine at every third amino acid in the triple helical domains of collagen. We found that all three major glycine tRNA isoacceptors were increased during chondrocyte and osteoblast differentiation, suggesting that acquisition of specialized tissue functions may include adjustment of the tRNA pools to sustain its characteristic protein output. We also found that tRNA supply increased in advance of codon demand along both chondrogenic and osteogenic trajectories, indicating a common differentiation program that upregulates collagen expression and tRNA biogenesis in parallel, instead of a program that upregulates tRNA expression through direct sensing of codon demand. The positive correlations between tRNA supply and codon demand changes in several musculoskeletal trajectories (chondrocyte, osteoblast, matrix fibroblast, myotube) further raised the possibility that the extent of tRNA remodeling was linked to the composition of specialized protein output. In contrast, weaker correspondence between tRNA supply and codon demand changes in neuronal trajectories indicated a relatively stable tRNA anticodon pool. A study demonstrated neuronal-specific sensitivity to depletion of particular tRNA isoacceptors (*56*). Therefore, selective tRNA isoacceptor requirements may coexist with relatively stable tRNA anticodon pools. Furthermore, amino acid-specific discordance between tRNA supply and amino acid demand can motivate investigations of whether tRNA charging, editing, modification, processing or turnover adjusts functional tRNA availability independently of tRNA abundance.

In CNS development, we identified regionally enriched lncRNAs and shared snoRNA dynamics, suggesting that ncRNA programs accompanied both neuronal differentiation and diversification across distinct neuronal identities. Predicted miRNA convergence on progenitor and gliogenic regulators raised the possibility that post-transcriptional repression reinforced neuronal commitment, although functional perturbations were needed to establish these regulatory relationships. Extending observations from human cell culture, the shared embryonic induction of *Snord116* across CNS populations provided a developmental context for the Prader–Willi syndrome and nominated E15–E16 as a window for testing how its loss affected neuronal maturation.

In summary, DT-MOCA delineated cell type- and lineage-specific ncRNA programs and identified ncRNAs associated with distinct developmental stages and lineage functions.

## Acknowledgments

We thank members of the Fan lab for helpful discussions.

## Author contributions

Conceptualization: RF

Methodology: HL, ML, RF, DZ, KL,

Investigation: ML, DZ, KL, YK, JL, MBG, RF

Visualization: ML

Funding acquisition: RF, ML

Project administration: HL, RF

Supervision: RF

Writing – original draft: ML

Writing – review & editing: ML, RF, HL

## Data, code, and materials availability

All instructions for preparing the materials or reagents are provided in the supplementary materials. No newly created materials are included in this study. The ASTRO pipeline used for raw data processing is downloaded from https://github.com/gersteinlab/ASTRO. Data analysis codes and processed data of the DT-MOCA atlas have been deposited to Dryad. The reference mouse prenatal development atlas encompassing 12.4 million nuclei is downloaded from https://omg.gs.washington.edu/. All raw sequencing data in this study will be deposited in the NCBI Gene Expression Omnibus before publication and made publicly available upon publication.

## Materials and Methods

### Single-nucleus total RNA-sequencing in FFPE tissue

#### Tissue deparaffinization and decrosslinking

Mouse embryo FFPE samples were obtained from Zyagen. Animal care and tissue collection were approved by the in-house Institutional Animal Care and Use Committee of Zyagen. FFPE sections were preprocessed with deparaffinization and decrosslinking. The samples were further rinsed with nuclease-free water and dried with compressed air, followed with permeabilization with permeabilization solution containing 0.5% Triton X-100 in 0.5X PBS-RI for 15 minutes at room temperature. The tissue was then rinsed with 0.5X PBS-RI followed by air dry.

#### In situ enzymatic reaction

For polyadenylation, poly(A) wash buffer comprised of 1X Poly(A) Polymerase Reaction Buffer, 0.6 U/µL Qiagen RNase Inhibitor and 0.6 U/µL SUPERase·In RNase Inhibitor was applied to the tissue for 5 minutes at room temperature. The tissue was washed briefly with water and air dried. In situ poly(A) tailing was performed by adding poly(A) reaction mix (1X Poly(A) Reaction Buffer, 1 mM ATP, SUPERase·In RNase Inhibitor, Qiagen RNase Inhibitor, NEB Poly(A) Polymerase) to the tissue, followed by a 37°C incubation for 30 minutes. The tissue was washed twice with 0.5X PBS-RI wash buffer. In situ reverse transcription was performed by applying the reverse transcription reaction mix comprising 1X Maxima RT buffer, biotin RT Primer, dNTPs, SUPERase·In RNase Inhibitor, Qiagen RNase Inhibitor, and Maxima H Minus Reverse Transcriptase onto the tissue section. The reaction was incubated inside a humidified chamber at room temperature for 30 minutes, followed by a 42°C incubation for 60 minutes. Next, template switching oligo reaction buffer (Template Switching Oligo, 1X Maxima RT buffer, Maxima H Minus Reverse Transcriptase) was directly added into the RT mix. Template switching reaction was performed inside a wet box at room temperature for 30 minutes and 42°C for 30 minutes. The tissue was then washed with 0.5X PBS-RI wash buffer, rinsed with water, and air dried completely.

#### Combinatorial split-pool barcoding and ligation

Following single nuclei isolation, the nuclear pellet was resuspended in the hybridization mix containing 1X T4 ligation buffer and nuclei isolation buffer. When multiple samples were processed simultaneously, the total resuspension volume was divided among samples in proportion to their nuclei counts. Combinatorial barcoding was carried out in three successive rounds of “split-pool”. In the first round, 40 µL of the nuclear suspension was added to each well of a 96-well plate. The plate was incubated at room temperature for 30 minutes under gentle shaking at 300 rpm. Unbound sites were blocked by adding 10 µL of the round 1 blocking solution to each well, followed by a 30-minute incubation at room temperature with gentle shaking. Nuclei from all 96 wells were pooled into a reagent reservoir, mixed thoroughly, and redistributed at 50 µL per well into a 96-well barcoding plate. The plate was incubated at room temperature for 30 minutes with gentle shaking. The round 2 blocking solution was added to each well with 10 µL per well and incubated for 30 minutes at room temperature with gentle shaking. For the third round, the nuclei were pooled into a new reagent reservoir and redistributed at 60 µL per well into a 96-well third-round barcoding plate. After a 30-minute incubation at room temperature with gentle shaking, 10 µL of the round 3 blocking solution was added to each well and incubated for an additional 30 minutes. Finally, barcoded nuclei from all wells were pooled together and centrifuged at 1000g for 4 minutes at 4°C to remove the supernatant. The nuclear pellet was washed twice with nuclei isolation buffer, with centrifugation at 1000g for 4 minutes at 4°C between washes. The nuclei pellet was resuspended in the ligation mix (1X T4 ligation buffer, 20 U/µL T4 DNA ligase, nuclear isolation buffer). The ligation reaction was incubated at room temperature for 20 minutes.

#### cDNA amplification and library preparation

Following ligation, nuclei were washed, lysed and decrosslinked using proteinase K, which was subsequently inactivated with PMSF. Biotinylated cDNA was captured on streptavidin beads and amplified by PCR, with additional cycles determined by quantitative PCR. Amplified cDNA was purified using SPRI beads and assessed by TapeStation. Two rounds of rRNA depletion were performed using the SEQuoia RiboDepletion Kit, followed by final library amplification and purification. Libraries were sequenced on an Illumina NovaSeq X Plus with 150-bp paired-end reads. A detailed summary of individual embryos can be found in **table S1**. All primer, linker, blocking oligo and barcode sequences were included in **table S2**.

No statistical methods were used to determine sample size. This study was designed to construct an atlas of coding and noncoding RNA expression during mouse organogenesis. Sampling prioritized coverage of developmental stages and cellular diversity. Twelve FFPE tissue sections spanning eight embryonic stages, E11–E18, were processed across three experimental runs to characterize cell-type-specific expression and developmental trajectories. The sample size enabled the identification of 21 major cell types with high concordance with an independent *Qiu et al.* atlas (*10*), revealed complex developmental trajectories of the musculoskeletal system and neuronal systems, and recovered significant CNS neuron cell types and brain regions. Embryos used in experiments were randomly selected from each timepoint before sample preparation. In this study, investigators were blinded to group allocation during sample acquisition, data collection, and data analysis. Embryo acquisition, data collection, and data analysis were performed by different researchers in different locations.

### snTotal-FFPE data preprocessing

Raw data processing uses the ASTRO pipeline (*54*). Single-cell gene-expression data from mouse embryos were analyzed in Python v3.12.13 using Scanpy v1.12.1 (*55*). Intronic and exonic features were merged with their corresponding gene entries by summing counts. Cells were retained if they contained at least 250 total counts and 200 detected genes, and genes detected in fewer than 20 cells were excluded. Doublet scores were calculated separately for each sample using Scanpy’s implementation of Scrublet (*56*), with an expected doublet rate of 5%, and cells with doublet scores >0.4 were removed. Counts were normalized to 5,000 per cell and log1p-transformed. Highly variable genes were selected using mean expression thresholds of 0.0125–3 and a minimum normalized dispersion of 0.5. Total counts and mitochondrial count percentage were regressed out, and expression values were scaled to unit variance and clipped at 10 before principal component analysis using the ARPACK solver. Batch integration was performed by experimental batches using Harmony (*57*) through harmonypy v0.0.10. A neighborhood graph was constructed using 20 neighbors and cosine distance across the first 30 Harmony-corrected principal components. UMAP embeddings were generated with a minimum distance of 0.2. Leiden clusters were manually assigned to 21 major cell types based on canonical marker gene expression. Protein-coding marker genes used for cell type annotations were included in **table S3**.

To identify cell type-specific ncRNAs, we first normalized each ncRNA transcript to the total transcript count in each cell, scaled to 10,000 counts per cell, and log1p-transformed to account for differences in sequencing depth. The ncRNAs detected in less than five cells were excluded. Differential expression analyses were performed for each ncRNA biotype, including lncRNA, miRNA, snRNA, snoRNA, and tRNA. For each ncRNA biotype, expression in an annotated cell type was compared with expression in all remaining cells using Wilcoxon ranksum tests. The ncRNAs with an adjusted p<0.05 and a positive log2 fold change were identified as candidate cell type-enriched markers. For tRNAs, counts from genomic loci belonging to the same isodecoder family were summed within each cell before normalization and differential expression analysis. Cell type-specific ncRNAs for each RNA biotype were included in **table S4**.

### Gene classification by developmental time and cell type expression dependence

Raw UMI counts were aggregated into pseudobulks by experiment, embryonic day and cell type annotations, with pseudobulks containing fewer than 20 nuclei excluded. Cell types were retained only if represented by at least six pseudobulks spanning at least three embryonic days, which removed mast cells and T cells. The library size of each pseudobulk was the total UMI count over all genes within each pseudobulk. Genes were retained if detected in more than 2% of retained pseudobulks and supported by at least 10 UMIs in total. Each gene was fitted separately using a negative-binomial generalized additive model with a log link in mgcv version 1.9-4 under R version 4.4.1. For gene g in pseudobulk i, the model was specified as:

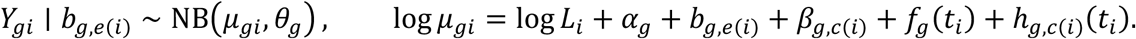

*L_i_* denotes pseudobulk library size, included as an offset; *α*_g_ is the gene-specific intercept; *b*_g,*e*(*i*)_ is an experiment-level random intercept; *β*_g,*c*(*i*)_ is a categorical cell-type term; and *t_i_* is embryonic day. The shared temporal component, *f*_g_(*t_i_*), was represented by a cubic regression spline with basis dimension five. The cell-type-specific component, ℎ_g,*c*(*i*)_(*t_i_*), was implemented using a sum-to-zero factor-smooth basis with cubic regression splines of basis dimension five (bs="sz", xt=list(bs="cr")). Corresponding spline coefficients were constrained to sum to zero across cell types, and each cell-type-specific smooth had a separate smoothing parameter. Benjamini– Hochberg adjustment was applied separately for each term across all modeled ncRNAs and, independently, across protein-coding genes. Significance thresholds were FDR <0.10 for miRNAs and <0.05 for all other gene types. Genes with a significant factor smooth term were assigned to the interactive class. Among the remaining genes, those with both significant main terms were classified as additive, those with only one significant main term as time-only or cell-type-only, and those with neither as not significant.

### Evolutionary age of lncRNAs across developmental expression classes

Evolutionary age annotations were obtained from the mouse lncRNA catalogue of *Sarropoulos et al.* (*34*) by matching gene identifiers or symbols. Genes without an exact match were mapped by overlap between annotated exons on the same genomic strand, using GENCODE vM25 and the published mouse annotation in the mm10 assembly. Coordinate-based matches required at least 1 bp of shared exonic sequence. When several catalogue loci overlapped a gene, the locus sharing the largest total number of exonic bases was selected. Exonic overlap was also used to resolve conflicting ages among exact matches. Within each lncRNA gene class, genes with resolved ages were counted as mouse-specific or assigned to approximately 20, 80, 90, 180 or 300 million year age categories. Unmapped genes, unresolved mapping conflicts, and catalogue entries labelled as “ambiguous” or “multimember family” were excluded from age distribution calculations. The evolutionary age and gene class of lncRNAs were included in **table S5**.

### Transcription factor binding at lncRNA promoters across developmental expression classes

For the transcription factor (TF) binding analysis, unresolved, antisense and divergent lncRNAs were excluded, together with loci failing opposite-strand protein-coding overlap checks. Promoters extended 2 kb upstream and 1 kb downstream of the transcription start site of the transcript with the greatest summed exon length. Chromosome- and length-matched intergenic controls were sampled after excluding annotated genes, promoters, RepeatMasker intervals and assembly gaps. Promoters and controls were converted from GRCm39 to mm10 using the UCSC mm39ToMm10 chain and rtracklayer v1.66.0 (*58*). TF-binding sites were quantified by strand-independent overlap with all 832 downloaded mouse GTRD v20.06 TF metacluster tracks, without TF-expression filtering (*59*), using GenomicRanges v1.58.0 (*60*). Within each region, identical metacluster coordinates assigned to the same TF identifier were counted once. Adjacent plotted groups were compared using two-sided Wilcoxon ranksum tests with continuity correction in R v4.4.2.

### Nomination and visualization of the candidate competing endogenous RNA (ceRNA) network

Cell-type-specific co-expression networks were constructed in R 4.4.1 using hdWGCNA 0.4.12, with WGCNA 1.72-5 providing the underlying network analysis framework and Seurat 5.5.1. Raw UMI counts were aligned with cell annotations and Harmony embeddings from the annotated h5ad object. In each cell type, protein-coding and lncRNA markers with adjusted P ≤ 0.05 and positive log fold change were retained for module construction. Metacells were generated separately within cell type and embryonic origin groups using nearest neighbors in the Harmony space, with an adaptive neighborhood size of up to 25 cells and maximum shared cell count of floor(k/2). Summed metacell counts were normalized to 10,000 counts using full-transcriptome library sizes and log1p-transformed. Signed networks were constructed using biweight midcorrelation and signed topological overlap. Modules were identified using a minimum module size of 30 genes, deepSplit = 4, and merging cut height of 0.2. Module eigengenes were represented by the first principal component, and gene–eigengene correlations quantified module membership, or kME. Soft-thresholding powers were selected separately for each cell type using a scale-free topology target of R² ≥ 0.8, a negative fitted slope, and mean connectivity ≥ 1. Representative subnetworks in **Fig. S4** contained up to 25 lncRNAs and 25 protein-coding genes per module, ranked by kME. Signed topological overlap values were reconstructed from saved metacell expression using the previously selected soft power and all valid saved network genes. Displayed edges included connections at or above the 92nd percentile of within-subnetwork topological overlap together with each node’s three strongest connections. Networks were arranged using a weighted Fruchterman–Reingold layout. Node area represented kME, while edge width and opacity represented topological overlap. Functional enrichment was performed on complete module gene sets using clusterProfiler 4.12.0 and ReactomePA 1.48.0, with detected genes in the corresponding cell type network serving as the background. GO Biological Process, Cellular Component and Molecular Function, KEGG, WikiPathways and Reactome terms were retained at Benjamini– Hochberg-adjusted P < 0.05, and up to two terms per annotation category were displayed for each selected module.

For ceRNA analysis, the same module assignments were used to identify lncRNA–mRNA pairs sharing miRNA interaction evidence from ENCORI/starBase and multiMiR. The miRNA loci were included in downstream correlation tests but were excluded from hdWGCNA module construction. Candidate triplets required positive lncRNA–mRNA and negative miRNA–mRNA Spearman correlations across log-normalized metacells, with nominal BH-adjusted P < 0.1 for both. In **Fig. 3**, miRNA–lncRNA correlations were negative with nominal adjusted P < 0.1. Candidates were further prioritized using miRNA detection, genomic-confound annotations, developmental expression coherence and functional enrichment. The selected triplets were displayed as networks containing lncRNA, miRNA and mRNA nodes, with uniformly thick solid edges colored by relationship type. These networks represent exploratory, interaction-supported ceRNA hypotheses because expression correlations do not establish competition.

### Developmental trajectory construction and ncRNA dynamics

Developmental trajectories were reconstructed using Slingshot v2.12.0 in R v4.4.1 using the precomputed 50-dimensional principal component representation and cell-type annotations. Noncoding RNA expression dynamics were modeled jointly across the inferred lineages using negative-binomial generalized additive models in tradeSeq v1.18.0, with five spline knots and the natural logarithm of total cellular library counts as an offset. Pseudotime-associated expression changes were assessed using lineage-specific associationTest tests, with P values adjusted across all modeled ncRNA features separately for each lineage using the Benjamini–Hochberg procedure. The lncRNA heatmaps included features with adjusted P <0.05; the other four ncRNA classes retained significant features or features detected in at least 10 cells retained for that lineage’s bins. Displayed features were required to have converged models and finite predictions. Cells with finite pseudotime and lineage weight ≥0.5 were ordered by pseudotime and divided into 15 bins containing equal numbers of cells. Fitted expression was predicted on a 200-point pseudotime grid and linearly interpolated to each bin’s mean pseudotime. Predictions were transformed as ln(1 + expression), standardized per feature across bins, and clipped to ±2.5 for display. Within each ncRNA class, significant features were grouped using iterative correlation clustering initialized with early, transient, increasing, decreasing, and late expression templates, then ordered by group and peak-expression pseudotime, with nonsignificant displayed features followed. The heatmap was generated using ComplexHeatmap v2.20.0, with proportional blocks for the five ncRNA classes and no additional hierarchical clustering. Columns were displayed at equal widths and annotated with scaled pseudotime and mean embryonic day. Selected literature-supported features were labeled using mouse gene symbols. In **table S6**, for each lineage, observed ncRNA expression was summarized across equal-cell pseudotime bins. Raw counts were summed within each bin and normalized to counts per million using the summed full-library counts of the constituent cells. Tables included feature identifiers and annotations describing each bin’s cell count, mean pseudotime and mean embryonic day.

### Amino acid codon demand and tRNA supply quantification

The amino acid codon demand and tRNA supply quantification methods in **Fig. 2** are similar to the method described in a prior publication(*30*). To estimate amino acid codon demand, we obtained mouse protein sequences from GENCODE M38 and selected the longest protein isoform for each protein-coding gene. Absolute counts of the 20 standard amino acids were tabulated for each selected sequence. Protein-coding features were matched to GENCODE by gene symbol, excluding ambiguous or unmatched mappings. Raw gene-level UMI counts were summed across all cells within each cell type or embryonic day. For each amino acid, demand was calculated by multiplying the summed UMI count of each gene by the corresponding amino acid count in its representative protein sequence and summing across all genes. This yielded a demand matrix with amino acids as rows and embryonic days as columns. To quantify tRNA supply, we summed raw UMI counts across tRNA features assigned to the same amino acid across cells within each cell type or embryonic day. Additionally, in **Fig. 5**, we analyzed tRNA supply and codon demand changes along developmental trajectories. Cells with finite Slingshot pseudotime and lineage weight ≥ 0.5 were ordered and divided into equal-cell bins. Raw UMI counts were summed within bins. Codon demand was calculated by multiplying each gene’s UMI count by codon counts in its representative Ensembl GRCm39 coding sequence and summing across mapped protein-coding genes. Amino-acid demand was obtained by summing synonymous codon demand and normalizing to total expression-weighted amino acid demand across all 20 amino acids in the bin.

For tRNA supply, the amino acid-specific tRNA supply was defined as tRNA UMIs assigned to the amino acid divided by total tRNA UMIs assigned across 20 standard amino acids. The **Fig. 5G** displayed the mean of the final two bins minus the mean of the first two bins for all amino acids across all reconstructed lineages.

## Supplementary Figures and Captions

**Fig. S1.**
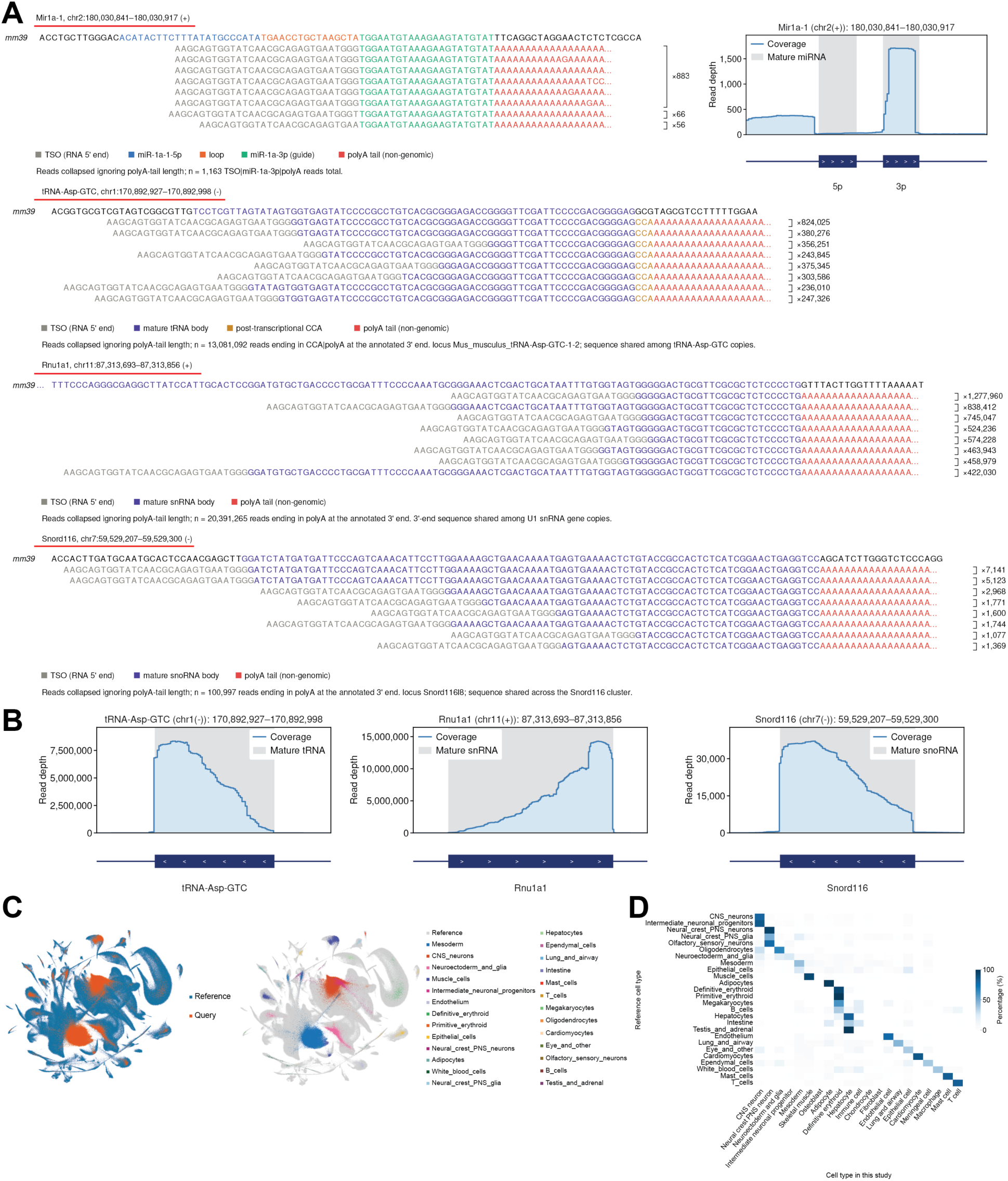
Performance benchmarking of ncRNA detection and atlas construction. (**A**) Representative raw sequencing reads aligned to mm39 reference sequences for *Mir1a-1*, *tRNA-Asp-GTC*, *Rnu1a1*, and the *Snord116* gene cluster. Numbers indicated read counts collapsed irrespective of the poly(A) tail length. The upper right panel showed the read coverage across the *Mir1a-1* locus, with mature 5p and 3p miRNA arms shown in gray. (**B**) Read coverage for *tRNA-Asp-GTC*, *Rnu1a1*, and the *Snord116* gene cluster, with mature regions shown in gray. (**C**) UMAP visualization of integrated reference and query datasets was displayed on the left. The same UMAP with nuclei colored by inferred query data cell types was displayed on the right. (**D**) Heatmap comparing cell type annotations in this study with reference cell type labels transferred using the Harmony-based k-nearest-neighbor classifier in popV (*61*). Colors indicated the percentage of nuclei assigned to each query cell type within each transferred reference category.

**Fig. S2.**
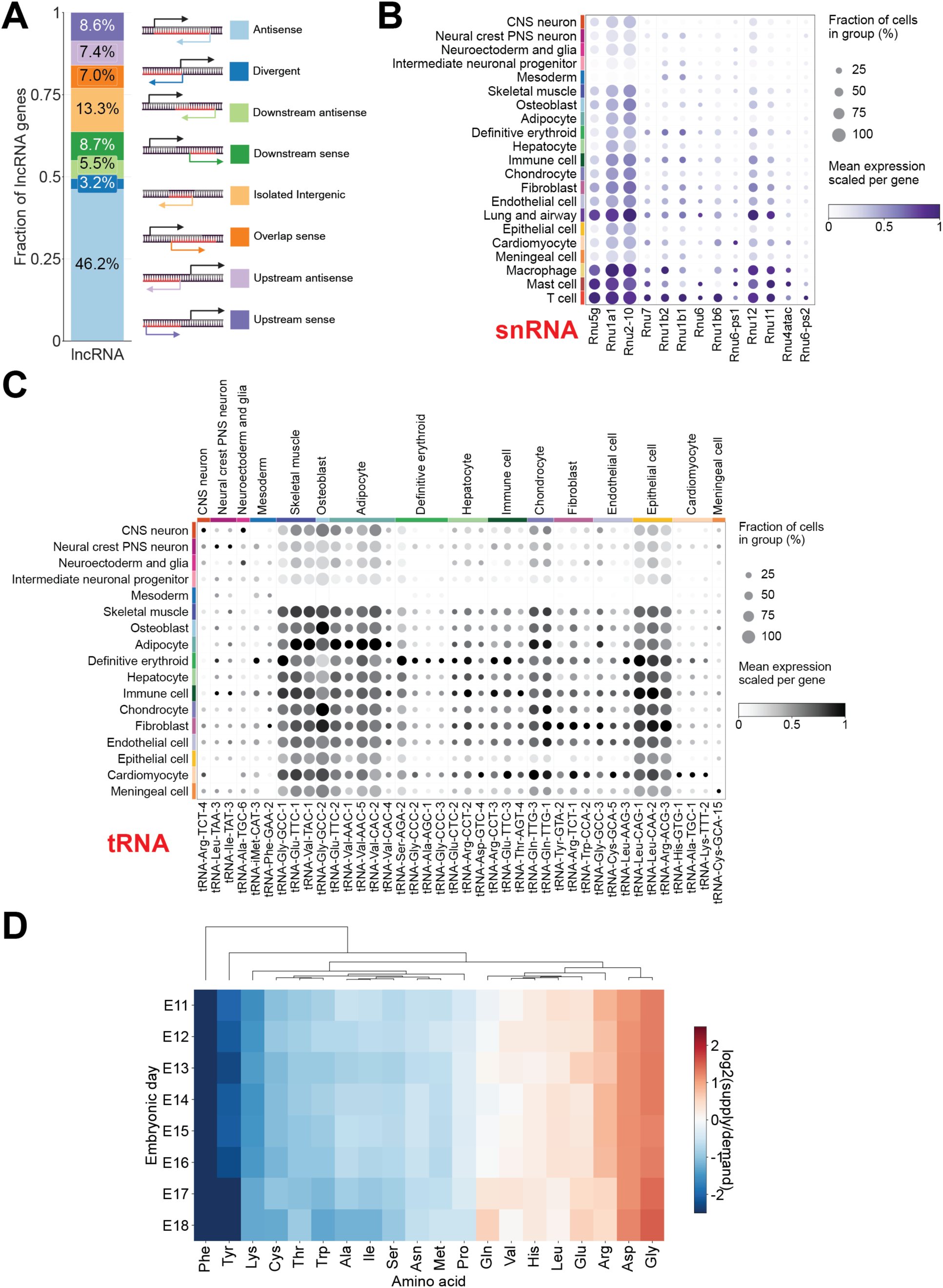
Cell type-specific ncRNA expression. (**A**) Fraction of lncRNAs in each genomic class. (**B**) Dot plot showing expression of representative snRNAs that were differentially expressed in each cell type. (**C**) Dot plot showing expression of representative tRNA isodecoders that were differentially expressed in each cell type. UMI counts were aggregated across genomic copies of each tRNA isodecoder. (**D**) Heatmap showing the ratio of tRNA supply to amino acid demand across embryonic days, with amino acids hierarchically clustered using Euclidean distance.

**Fig. S3.**
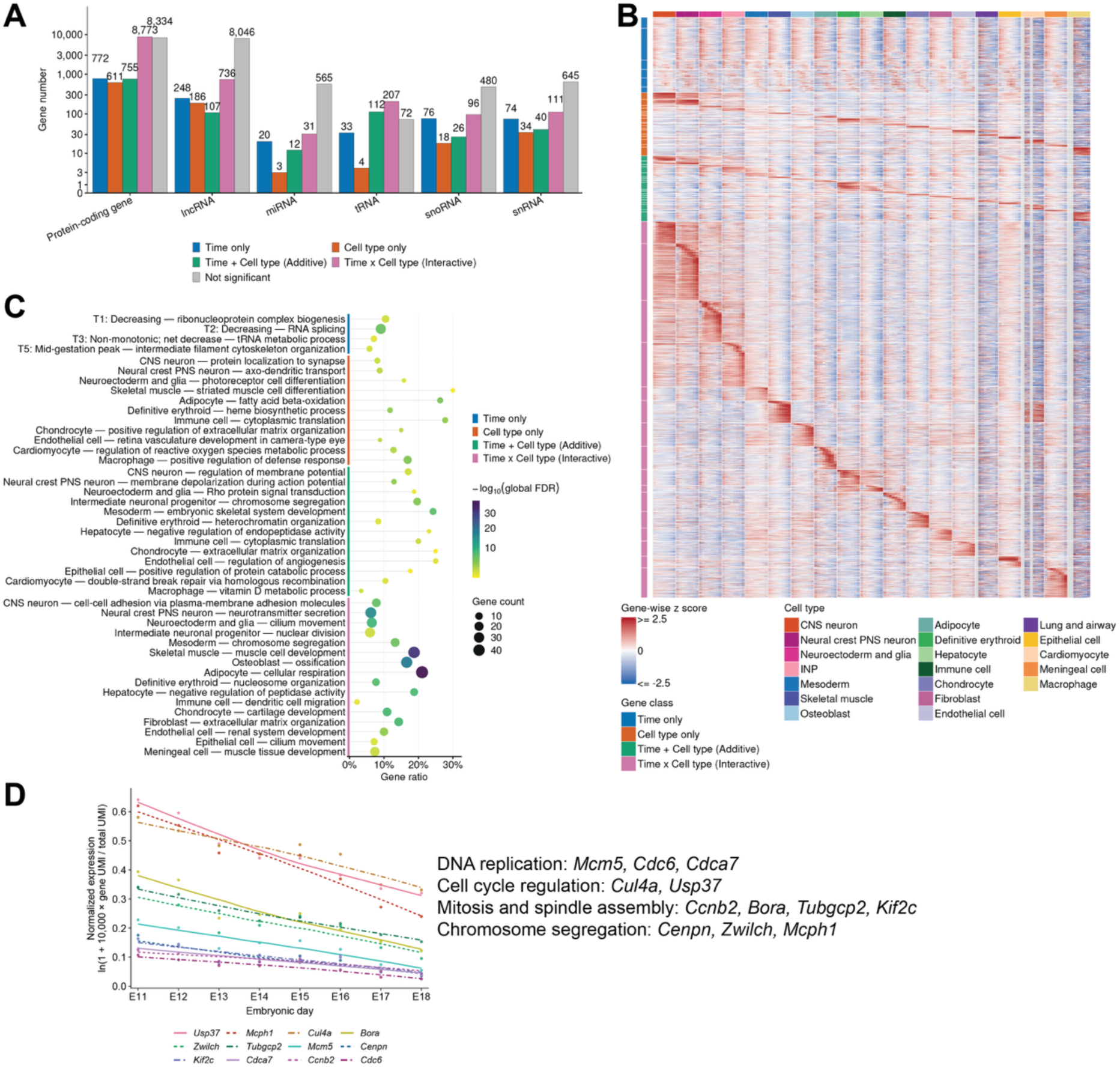
Genes fitted with the generalized additive regression model. (**A**) Gene number of genes assigned to each gene class for each RNA biotype. (**B**) Heatmap of model-fitted expression for protein-coding genes, with rows grouped by the four gene classes and expression pattern modules and columns ordered from E11 to E18 within each major cell type. Colors represented gene-wise z-scores of log2(fitted CPM+1). CPM: counts per million. Gray indicated cell type–day combinations lacking retained pseudobulk observations. (**C**) Gene ontology biological process enrichment of protein-coding expression modules grouped by gene classes, showing one representative enriched term per module. (**D**) Gene expression of selected proliferation and cell cycle genes classified as the Time-only class with decreasing expression from E11 to E18.

**Fig. S4.**
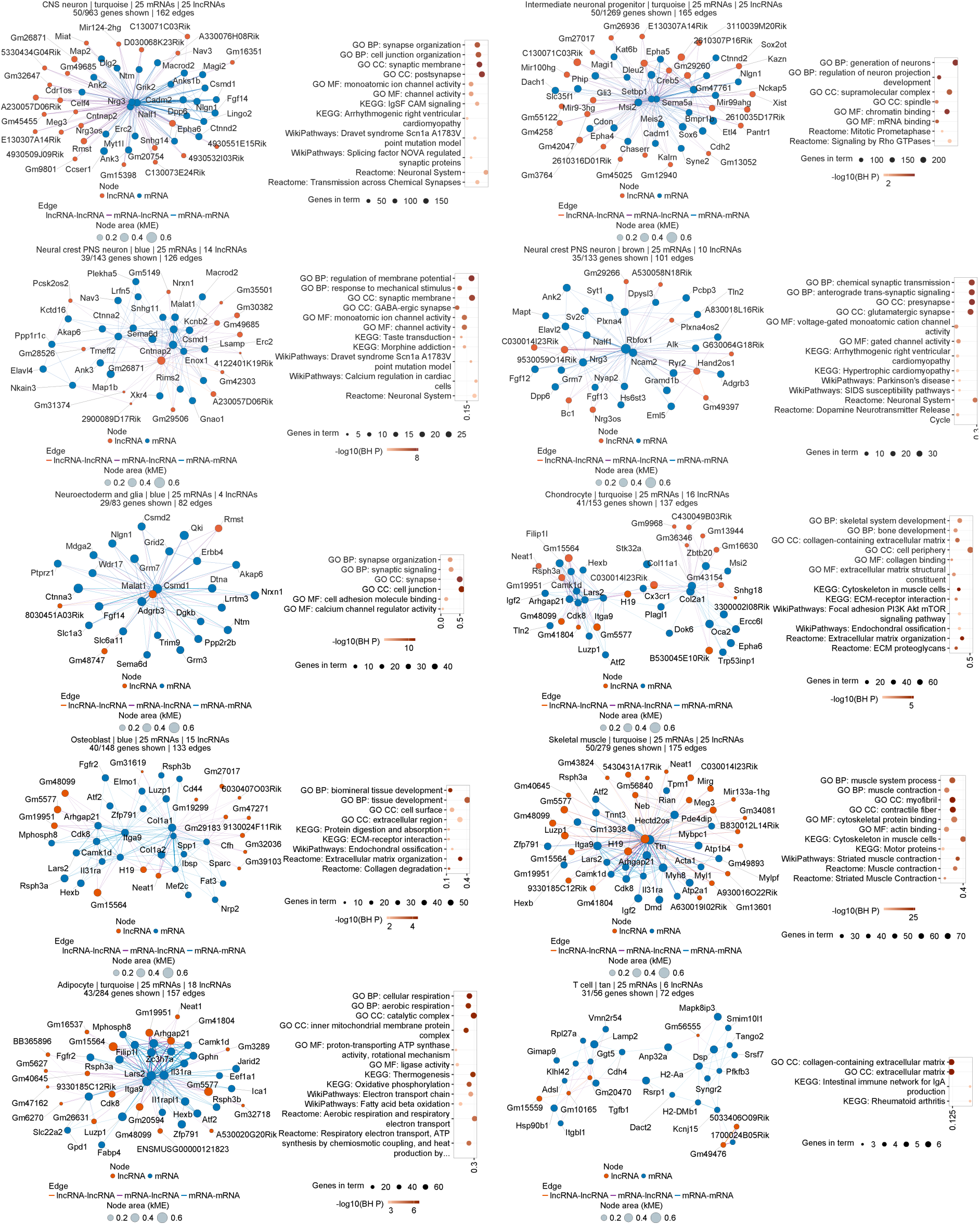
Gene co-expression networks containing both lncRNAs and mRNAs in each cell type. Selected hdWGCNA co-expression networks showing mRNAs (blue) and lncRNAs (orange), with node area indicating module membership (kME) and edge width reflecting topological overlap. Adjacent dot plots showed enriched Gene Ontology and pathway terms for each module.

**Fig. S5.**
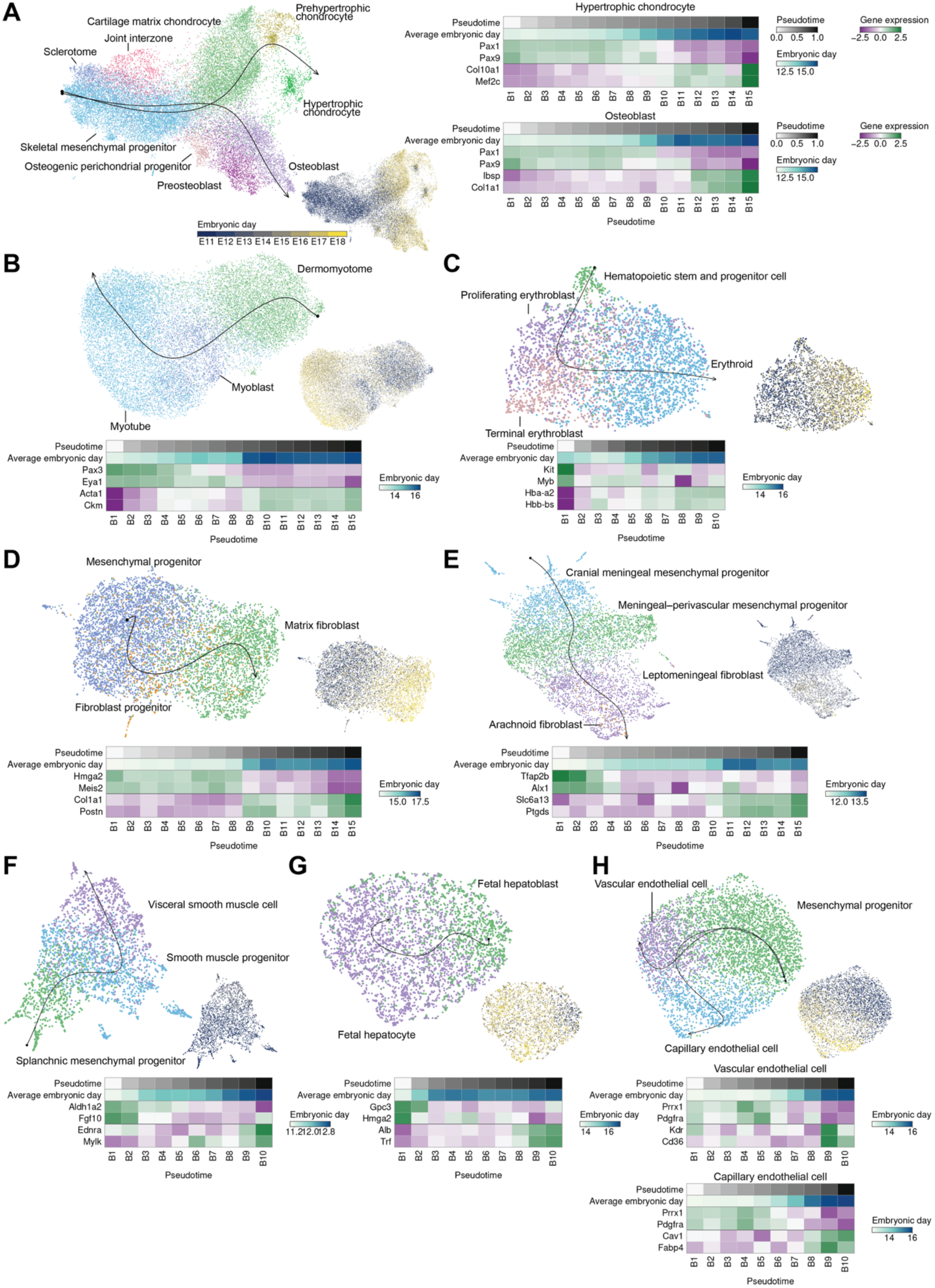
Characterization of protein-coding gene expression dynamics along non-neuronal developmental trajectories. UMAP of single-nucleus total RNA expression, with nuclei colored by cell types during the development of the skeletal system (**A**), skeletal muscle (**B**), hematopoiesis (**C**), fibroblast (**D**), meningeal cell (**E**), smooth muscle (**F**), fetal hepatocyte (**G**), and endothelial cell (**H**). Developmental trajectories fitted from Slingshot were projected onto the UMAP plots. The same UMAP with nuclei colored by embryonic days was displayed on the lower right of each panel. The heatmap showing expression of early-stage and late-stage marker genes along each trajectory was shown in each panel. Colors indicated gene-wise z-scores of gene expression. Upper bars indicated pseudotime and average embryonic day.

**Fig. S6.**
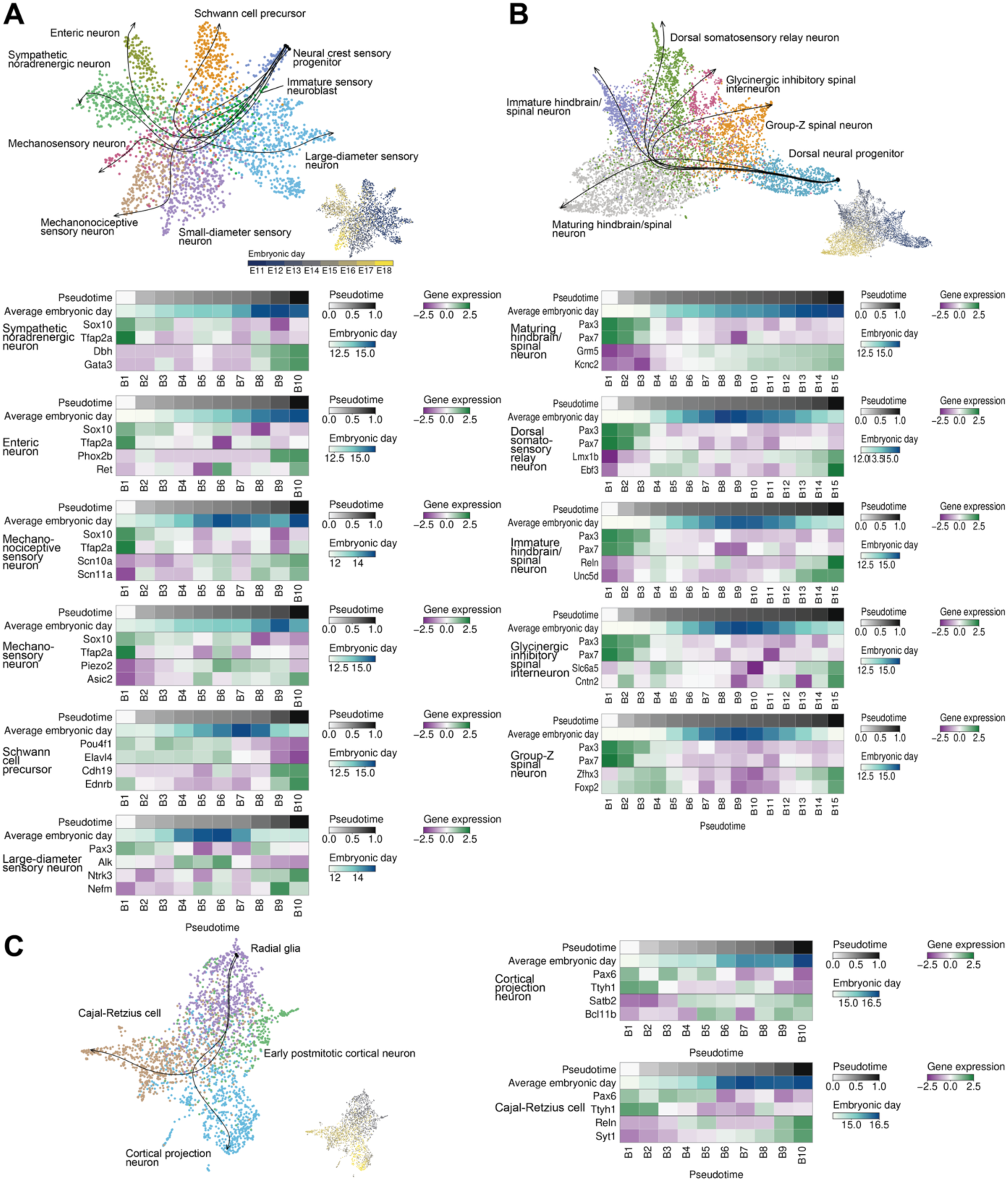
Characterization of protein-coding gene expression dynamics along neuronal developmental trajectories. UMAP of single-nucleus total RNA expression, with nuclei colored by cell types during the development of the neural crest (**A**), dorsal hindbrain/spinal neuronal system (**B**), and cortical neuronal system (**C**). Developmental trajectories from Slingshot were projected onto the UMAP plots. The same UMAP with nuclei colored by embryonic days was displayed on the lower right of each panel. The heatmap showing expression of early-stage and late-stage marker genes along each trajectory was shown in each panel. Colors indicated gene-wise z-scores of gene expression. Upper bars indicated pseudotime and average embryonic day.

**Fig. S7.**
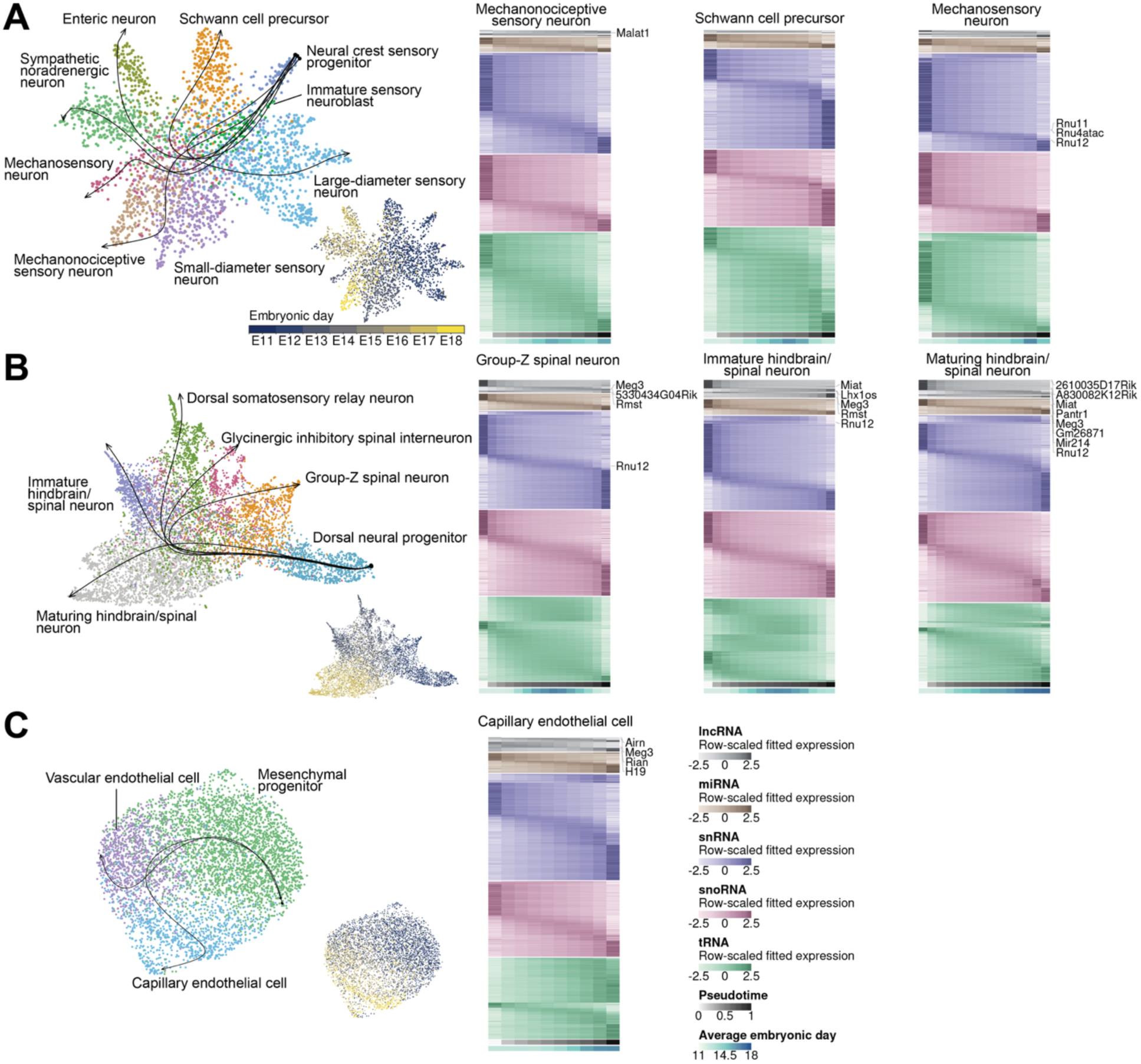
Characterization of ncRNA gene expression dynamics along developmental trajectories of the neural crest, dorsal hindbrain/spinal neuronal system and endothelial cell. UMAP of single-nucleus total RNA expression, with nuclei colored by cell types during the development of the neural crest (A), dorsal hindbrain/spinal neuronal system (B), and cortical neuronal system (C). Developmental trajectories fitted from Slingshot were projected onto UMAP. Heatmaps showed expression dynamics for lncRNAs, miRNAs, snRNAs, snoRNAs, and tRNAs along developmental trajectories, with rows grouped by RNA biotype and temporal expression patterns and columns representing equal-cell pseudotime bins. Colors indicated gene-wise z-scores of gene expression. Bottom bars indicated pseudotime and mean embryonic day.

**Fig. S8.**
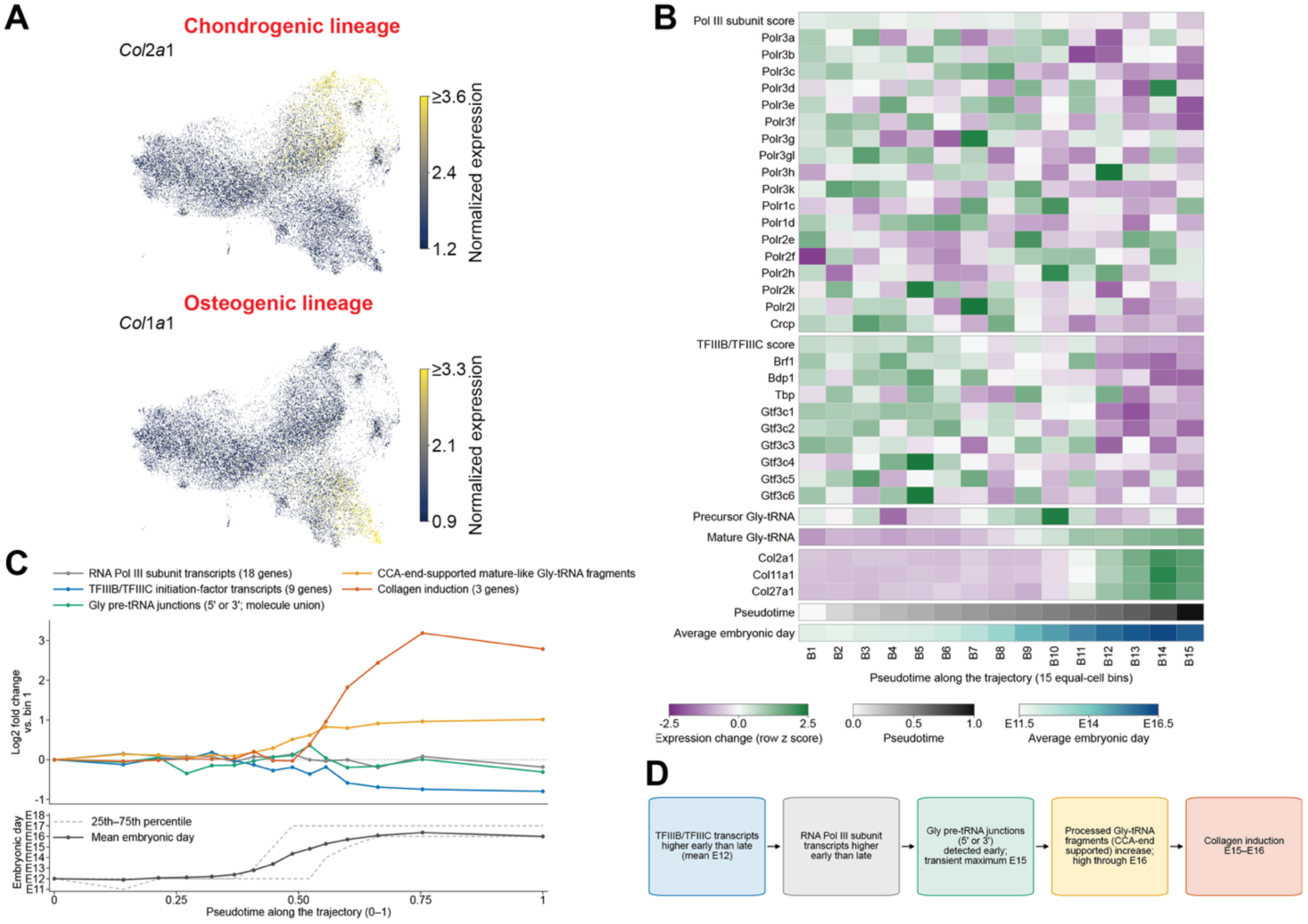
Pol III machinery, Glycine tRNA expression and collagen gene expression during skeletal development. (**A**) UMAP plot showing *Col2a1* and *Col1a1* expression along the chondrogenic and osteogenic lineages. (**B**) Heatmap showing log-normalized expression z scores of Pol III machinery genes, Glycine tRNAs, and collagen genes across equal-cell bins along the chondrogenic pseudotime. Pol III and TFIIIB/TFIIIC scores averaged constituent gene z scores. Precursor Gly-tRNAs were identified by leader/body or body/trailer junctions. Mature-like fragments were identified by processed CC/CCA-supported ends. Bottom tracks showed pseudotime and average embryonic day. (**C**) Relative expression changes along chondrogenic pseudotime. Curves showed relative expression of genes of the Pol III subunits, TFIIIB/TFIIIC factors, precursor Gly-tRNAs, mature-like Gly-tRNA fragments and collagens (*Col2a1*, *Col11a1*, *Col27a1*) across pseudotime bins. Summed counts were normalized to total UMIs and expressed as log2 fold change relative to that of bin 1. The lower panel showed mean embryonic day (black) and the interquartile boundaries (gray dashed lines). (**D**) Schematic summarizing temporal gene expression patterns during chondrogenic differentiation.

**Fig. S9.**
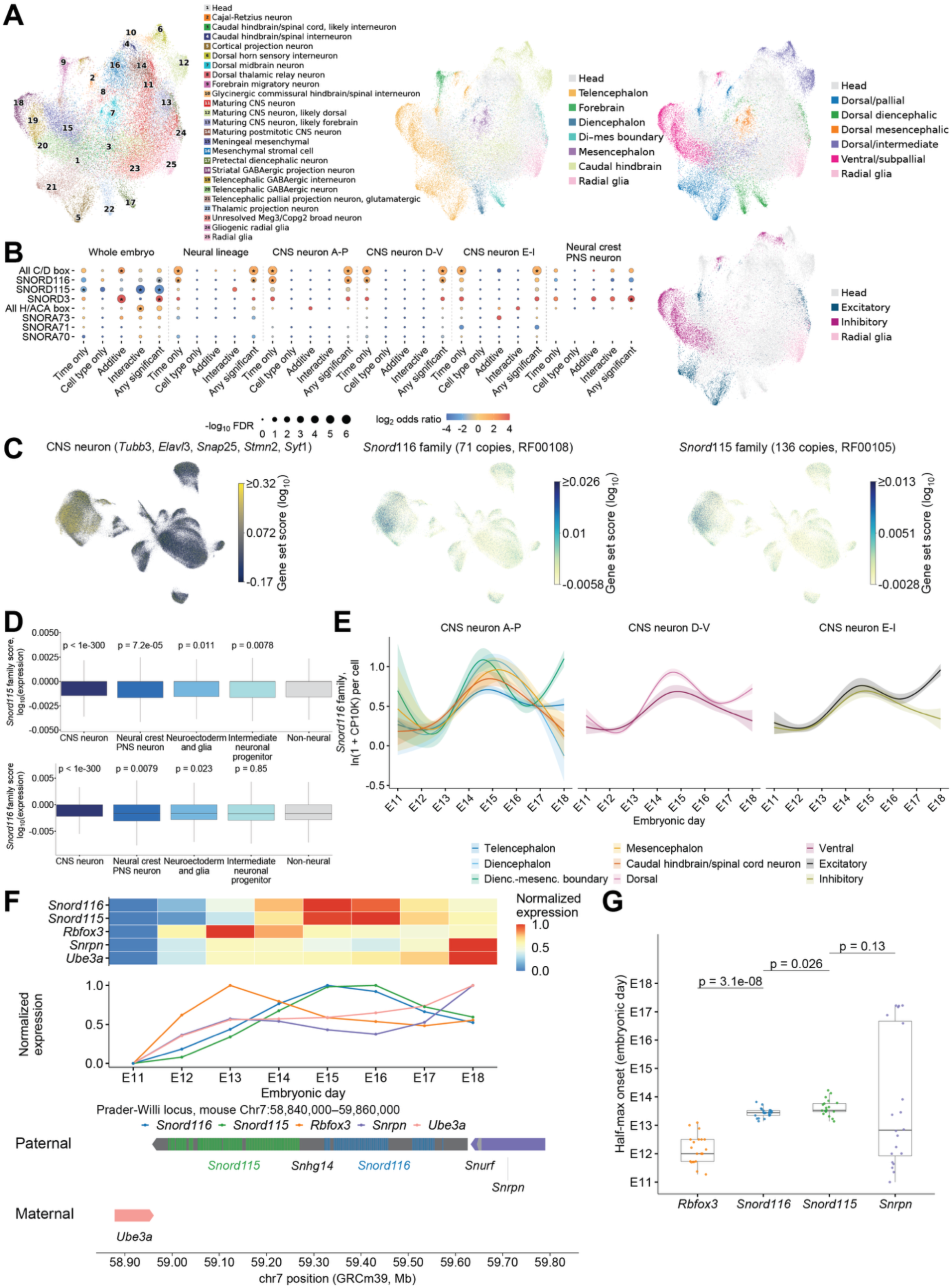
Developmental and cell type-specific expression of snoRNA genes during mouse embryogenesis. **(A)** UMAP plots showing single-nucleus total RNA expression in CNS neurons, with nuclei colored by cell type, anterior-posterior position, dorsal-ventral position, or excitatory/inhibitory identity. (**B**) Enrichment of snoRNA family genes in each gene class derived from generalized additive regression models across the whole embryo, neural lineage, CNS anterior-posterior group, CNS dorsal-ventral group, CNS neurons with excitatory and inhibitory functions, and neural crest lineage. (**C**) UMAP plots showing the CNS neuron (*Tubb3, Elavl3, Snap25, Stmn2, Syt1*) marker gene score, *Snord116* family gene score, and *Snord115* family gene score. (**D**) Distribution of *Snord115* and *Snord116* family gene scores across neural cell types and pooled non-neural cells. Boxes indicated the interquartile range, with horizontal lines marking the median and whiskers extending to the most extreme observations within 1.5× the interquartile range of box boundaries. *P* values were calculated with two-sided Wilcoxon ranksum tests comparing each neural cell type with non-neural cells. (**E**) *Snord116* family gene expression across E11-E18 in CNS neuron groups defined by anterior-posterior position, dorsal-ventral position, and excitatory/inhibitory identity. *Snord116* family UMIs were summed per cell, divided by total cellular UMIs, scaled to 10,000 and transformed as ln(1+normalized counts). Curves were fitted using geom_smooth, with shading showing 95% confidence intervals. (**F**) Expression of *Snord116*, *Snord115*, *Rbfox3*, *Snrpn* and *Ube3a* across E11–E18, averaged across neural cell types and scaled to 0–1 for each gene or family. (**G**) Distribution of estimated time when fitted expression first reached the midpoint of its developmental range for *Rbfox3*, *Snord116*, *Snord115* and *Snrpn*. Points represented neural cell types. Boxes indicated the interquartile range, with horizontal lines marking the median and whiskers extending to the most extreme observations within 1.5× the interquartile range of box boundaries. *P* values were calculated with two-sided Wilcoxon ranksum tests.

## Supplementary Tables

**Table S1.** Summary of individual embryo samples.

**Table S2.** Oligonucleotide sequences of primers, linkers, blocking oligos, and barcodes used for the DT-MOCA atlas construction.

**Table S3.** Protein-coding marker genes used for cell type annotations.

**Table S4.** Cell type-specific ncRNAs that are differentially expressed in each cell type.

**Table S5.** The evolutionary age and gene class of lncRNAs.

**Table S6.** Observed ncRNA expression across equal-cell pseudotime bins in each lineage. Raw counts were summed within each bin and normalized to counts per million using total RNA counts of the constituent cells. The table also included feature identifiers and the cell count, mean pseudotime and mean embryonic day of each bin.

